# Out-of-balance Growth Enables Cost-free Synthesis of the Flagellum and Other Proteins in a Single Bacterium

**DOI:** 10.1101/2024.12.17.629032

**Authors:** Mayra Garcia-Alcala, Josiah C. Kratz, Philippe Cluzel

## Abstract

The cost of expressing unnecessary genes on growth has been explained at the population level through balanced growth assumptions. However, due to the high degree of stochasticity in growth and gene regulation, it remains unclear how classical growth laws derived from bulk measurements manifest at the level of individual cells. For example, flagellar gene expression, one of the most energy-intensive processes in E. coli, is typically associated with reduced growth rates in bulk measurements. By contrast, we find at the single-cell level that a strikingly opposite behavior coexists: flagellar gene activity is associated with an increase in growth rate. We show that this apparent contradiction, reminiscent of Simpson’s paradox, results from examining in each cell instantaneous growth and gene expression without indiscriminately aggregating the single-cell data like in bulk measurements. We attribute this unrecognized behavior to out-of-balance growth boosts that temporarily offset the burden of flagellum production, likely driven by a surplus of growth factors inherited across cell division. Furthermore, we show that this non-intuitive relationship between growth and gene expression at the single-cell level is general, as we observe similar effects with synthetic systems driving the expression of fluorescent proteins downstream of constitutive promoters. Finally, we use a computational model to show that the inheritance of parental growth factors can explain the apparent contradiction and constitutes a general mechanism for mitigating the short-term cost of nonessential and biosynthetically demanding genes in the single cell.

## Introduction

The balance between the cost associated with the expression of genes and the demands for intracellular resources driven by growth is a tightly regulated process in bacteria, reflecting an evolutionary pressure to optimize fitness [1–4]. Recent studies have built on this principle to derive powerful, predictive growth laws [2, 5–7]. For example, at the population level, these laws predict that expressing unnecessary proteins reduces the availability of ribosomes that would otherwise be destined for the synthesis of other proteins, including those involved in ribosomal function and other cellular processes [6]. Consequently, the synthesis of unnecessary proteins is accompanied by a growth burden that increases with protein expression levels [1, 4, 7]. These laws have been derived under the assumption of balanced growth: a state where all cellular components, such as proteins, RNAs, DNA, and lipids, are synthesized at rates that are proportional to the constant growth rate of the cells measured at the population level [8, 9]. Balanced growth also ensures that cell growth has reached a steady-state and does not run into resource bottlenecks that could impair its physiology [10, 11]. While balanced growth is a practical assumption for predicting bacterial growth at the population level, it does not take into account out-of-balance processes [12–14], such as sudden and stochastic molecular events that inherently accompany growth and gene expression observed in single cells [15–22], even when environmental conditions are well-controlled and steady. Therefore, extracting the laws that govern the interplay between stochastic growth variations and protein synthesis is crucial for predicting and understanding the behavior and constraints of the single cell.

In recent years, studies in *E. coli* have reported that flagellar genes, responsible for the synthesis of one of the most energy-intensive bacterial organelles, exhibit large stochastic expression pulses [20]. The assembly of this complex organelle requires more than 30 different proteins that mostly constitute the basal body of a rotary motor, as well as a 20,000 subunit-long filament [23, 24] (Fig. S1A). During flagella synthesis, the cell directs a sizable portion of protein synthesis resources to this process [25–27]. While it has been demonstrated from bulk measurements that during balanced growth, the expression of this locomotion apparatus is regulated to compensate on average for changes in cell size across distinct growth conditions [28–30], far less is known about out-of-balance processes and their effect on gene expression, such as stochastic fluctuations of growth observed at the single-cell level. In view of the large protein synthesis demand that accompanies the stochastic transcriptional expression of flagellar genes, here we investigate the interplay between such gene activity and growth fluctuations within a single bacterium. Subsequently, we extend our analysis to that of synthetic expression systems and develop a computational model to validate the generality of our observations.

Collectively, production of flagellar proteins represents a sizable proteome investment that may consume resources otherwise available for growth, presenting a potential resource allocation tradeoff which the cell must navigate [30, 31]. The activity of flagellar genes follows a well-characterized transcriptional cascade [23, 32]. Briefly, it begins with Class-1 genes that encode the master regulator FlhDC, which drives the transcriptional activation of Class-2 genes. These genes are involved in the synthesis of the flagellar basal body, the hook, and the alternative sigma factor, FliA. This sigma factor, in turn, activates Class-3 genes that encode the filament and signaling chemotaxis proteins (Fig. S1A). Recently, the transcriptional activity of Class-2 and Class-3 flagellar genes has been found to be well-delimited in time with alternating and distinct *on* and *off* states, each spanning several cellular divisions [20]. Operationally, this temporal pattern makes the system experimentally accessible for dissecting the relationship between growth fluctuations and gene activity in living individual cells.

## Results

We tracked single cells as they grew and divided inside a microfluidic device called a ‘Mother Machine’ [33, 34], which allows bacteria to grow exponentially under controlled conditions for extended periods of time (Fig. S1B). This device confines cells within narrow channels, with one end exposed to a continuous flow of fresh medium while the other end is sealed. The temporal activity of flagellar promoters and the growth of individual cells are simultaneously monitored using fluorescence and phase-contrast microscopy (Materials and Methods). This experimental platform enables us to capture time-lapse images and extract various physiological parameters such as cell size, elongation rate, and fluorescence signals from reporter proteins with 5 min resolution over ∼30 generations (Fig. S1C). One key feature of this approach is its ability to isolate intrinsic fluctuations within the cellular system from those typically caused by uncontrolled environmental variations.

As previously observed by some of us, in the commonly used wild-type *E. coli* MG1655 strain, Class-2 and Class-3 promoters exhibit sparse stochastic pulses, whereas Class-1 displays low but sustained activity that resembles that of a constitutive promoter. The question then becomes how the growth rate varies during distinct periods of active or inactive flagellar gene activity. Here we use the elongation rate of cell size as monitored in the Mother Machine channels, as a measure of growth rate (Fig. S2). All experiments were performed in an MG1655 background harboring a *motA* mutation that disables swimming motility but does not impair flagellar assembly [35, 36], thereby isolating the regulatory and growth-related dynamics of the flagellar system (see Materials and Methods, Fig. S3). We monitored the activity of the Class-2 promoter, *fliFp*, using the fluorescent reporter SCFP3A as it also encapsulates the dynamics of Class-3, as previously described [20]. Practically, to capture the relationship between instantaneous variations in elongation rate and those of Class-2 activity, we extracted these two quantities from time series at each time point using a time window Δt = 5 min time window (Fig. 1 Left). These pairs were then sorted and grouped into bins based on their activity levels (Fig. S4). In addition to the strain with wild-type flagellar expression, we tested three distinct strains whose master regulator, FlhDC, is driven by unregulated synthetic promoters (P1, P2, P3; [20, 37]) with increasing strength, resulting in progressively higher transcriptional activity of downstream flagellar genes.

**Fig. 1.**
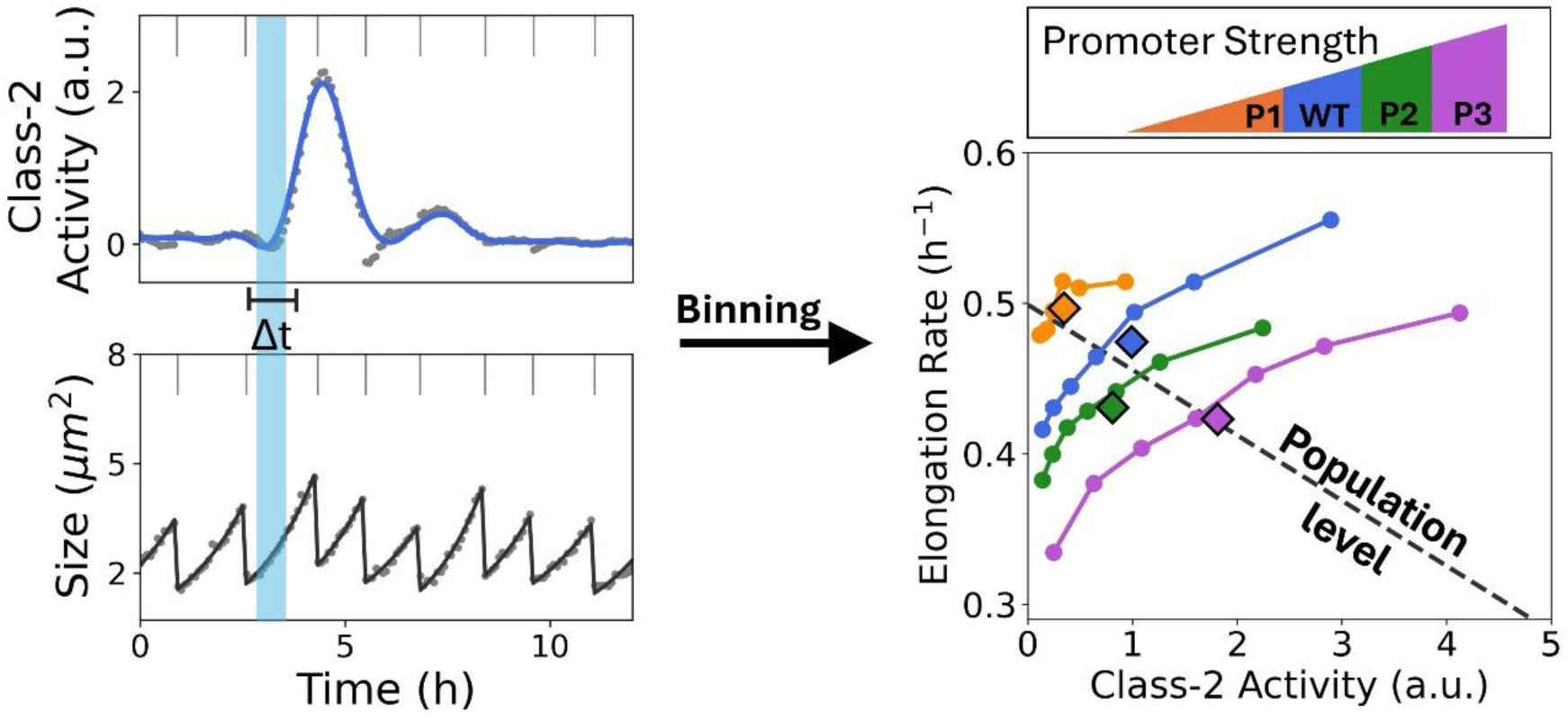
Relationship between instantaneous growth rate and flagellar gene activity in single cells versus population. Left: Time traces of Class-2 activity and Size (S) measured from the same mother cell across divisions (vertical ticks). We defined Class-2 activity as the time derivative of total fluorescence normalized by size, and the elongation rate as *ER*(*t*) = (1/*S*)(*ΔS*/*Δt*). Direct measurements of activity and size are shown as dots, whereas lines show the exponential fit for the size and the smoothed activity (see Methods and Materials). First, we used Class-2 activity to sort data points with Δt = 5 min and then bin the pair activity-elongation rate (see Fig. S4). Finally, we average the data in each bin and display the result in the right panel. Because the averaged bins are built from pairs of elongation rate values using a 5-minute time window, this binning allows us to determine the relationship between instantaneous flagellar gene activity and growth rate in single cells. Right: Instantaneous Class-2 activity versus elongation rate for different strains with varying mean levels of flagellar expression. In such strains, the native Class-1 promoter was replaced by synthetic promoters with different strengths, P1, P2 and P3 (color coded in top panel), P3 being threefold stronger than P1 [37]. The dashed line shows the linear fit to the population average of the pair Class-2 activity-elongation rate across strains, which illustrates the well-established relationship with a negative slope (cost) between growth rate and increasing mean flagellar gene activity across strains [30]. By contrast, single-cell binned data within each distinct strain shows a positive slope between instantaneous growth rate and flagellar activity, demonstrating that cells with higher flagellar gene activity are also the ones growing faster. Only the data exhibiting Class-2 activity (excluding *off-*states) are considered in this plot; see Fig. S3D for all states. The number of time traces is n= 69, 107, 96, 67 for the strains with the promoter P1, WT, P2, and P3, respectively, all with a duration of ∼47 hours.

The well-known population result is recovered when we fit the whole bulk of the instantaneous single-cell data across the four different strains (Fig. 1 Right, dashed line, Fig. S5), indicating that as the population of cells increases expression of unnecessary proteins, their growth rate decreases. Surprisingly, when investigating growth relationships for individual cells within each strain, we found that bins with higher instantaneous transcriptional activity were also associated with higher elongation rates (Fig. 1 Right, Fig. S5). This result is unexpected under the canonical growth laws and directly contrasts with population-level observations, for which elevated flagellar gene activity correlates with reduced growth rates (Fig. S6, [28–30]; see Fig. S7 for an alternative growth rate proxy). The coexistence of these opposing trends, positive within strains but negative across strains, suggests that a phenomenon such as the Simpson’s paradox is at play.

Intuitively, the Simpson’s paradox, a classic statistical phenomenon, can arise when distinct groups (i.e., individual strain) occupy different regions of a shared X–Y parameter plane (here, expression-growth). For example, each individual group might exhibit a positive internal trend between X and Y, but their respective data clusters are down shifted relative to one another. However, when we analyze the data across groups, this internal covariation within each group is masked. As a result, the aggregate analysis reflects only the relationship between these shifted group averages, completely reversing the underlying trend observed within the individual groups.

In our study, although single-cell and population analyses rely on instantaneous single-cell measurements, they differ fundamentally in how the data are aggregated, a distinction from which the observed paradox arises. In the population-level analysis (Fig. 1, dashed line), each point represents the mean instantaneous growth rate and mean promoter activity averaged across each distinct strain. This aggregation collapses co-fluctuations of growth and activity within cells and compares averages across genetic backgrounds, thus reflecting differences across strains governed by balanced growth constraints. In the binned analysis, each data point reflects paired fluctuations of growth and gene expression measured within the same cells, preserving their natural co-variation at a given time point. Hence, the binned analysis reveals dynamic coupling between growth and expression within single cells, whereas the aggregated data captures only the average trade-off observed across strains.

We performed the same analysis using the Class-1 promoter activity. In this case, the within-strain relationship between instantaneous promoter activity and elongation rate remained positive, but the trend was noticeably weaker than Class-2 (Fig. S8). We interpret this reduction as a consequence of Class-1 being noisy and that only a smaller fraction of Class-1 activity yields a productive Class-2. This behavior was explained in [20]. Thus, fluctuations in Class-2 activity more directly report changes in global translational capacity than Class-1 and therefore show a stronger coupling to growth.

We reasoned that if this single-cell effect reflects dynamic coupling between growth and Class-2 activity, it should also be observable along individual time series and not only in binned data. To examine this point further, we took advantage of the fact that flagellar genes show pulses of activity, which allow us to clearly track how growth rate changes during significant fluctuations in gene activity. We isolated time series segments of Class-2 activity featuring two consecutive pulses, rescaled their total duration to a unit interval (arbitrary units), and computed the average across a large collection of such pulse-to-pulse excerpts (Fig. 2, top; Materials and Methods).

**Fig. 2.**
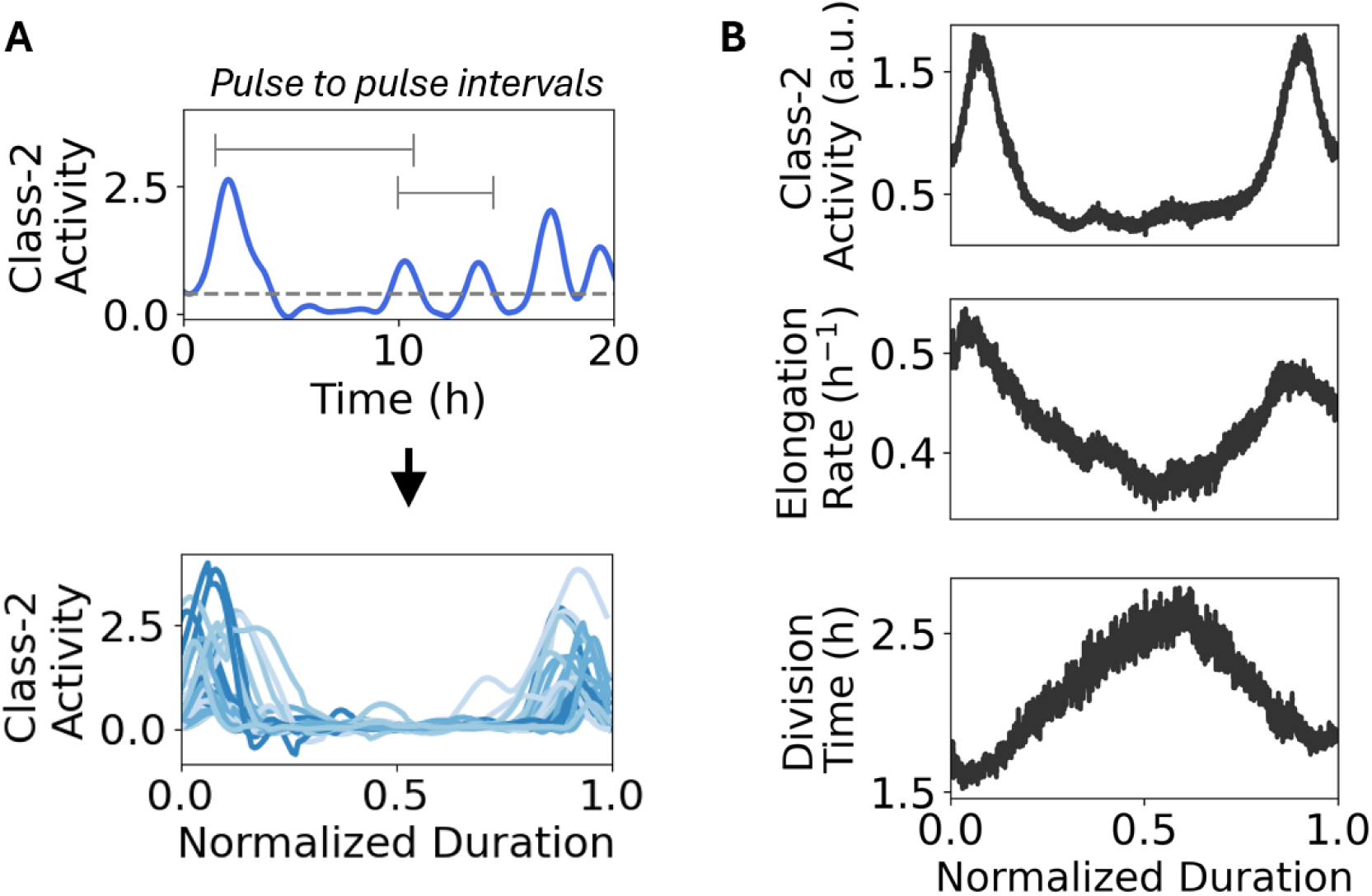
Cells grow faster during pulses. (**A**) The Class-2 activity time series were divided into intervals that consisted of a pulse of activity followed by an *off* state and another pulse. We selected all the intervals with similar durations (within a range of no more than 6 hours). The pulses were defined by the period for which the activity is above an empirical threshold (activity = 50 arbitrary units). Then, the intervals of similar duration were normalized such that they spanned from 0 to 1, in order to average them. (**B**) Top: Averaged pulse-to-pulse intervals (real time rescaled to 1) of the Class-2 activity. Middle and bottom: elongation rate and division time associated with the same time intervals as the top panel. The averages were calculated by applying a sliding window mean over the overlapping data consisting of n = 103 intervals.

Then, we jointly plotted the time series of elongation rate and division time that are directly associated with the pulse–to–pulse excerpts. We found that the aggregated time traces indicate that increases in flagellar gene activity coincide with higher elongation rates, while division time varies in the opposite phase (Fig. 2, middle and bottom). This relationship remains consistent across pulse-to-pulse intervals of different lengths (see Fig. S9).

To further examine whether there is a phase delay between elongation rate and Class-2 activity, we calculated the cross-correlation function of the full time series (about 20 divisions), to quantify how strongly elongation rate and activity are related when the activity signal is shifted in time (by a lag) relative to elongation rate. Thus, positive delays indicate that elongation rate lags behind promoter activity, whereas negative delays indicate that elongation rate precedes promoter activity. We found that the fluctuations in elongation rate temporally precede those of flagellar gene activity, and that they are positively correlated (Fig. S10, see Materials and Methods). This result was unexpected, since one might intuitively assume, based on population measurements, that increased flagellar gene activity would cause a cost that reduces the elongation rate. If that were the case, we would expect to see a negative correlation where changes in gene activity preceded those of elongation rate, rather than the positive correlation and temporal order we observed.

We repeated the correlation analysis using a reporter for the Class-3 promoter, *fliCp*, which drives expression of the energetically costly flagellin protein. We again observed a positive correlation (Fig. S11), with a slightly larger shift between elongation rate and activity for Class-3 compared to Class-2, consistent with Class-2 activity preceding and driving Class-3.

We also analyzed the three strains whose synthetic constitutive promoters (P1, P2, P3) control the expression of the flagellum master regulator and found similar temporal delay (Fig. S10). Since the constitutive promoters P1, P2 and P3 are synthetic, they do not respond to any input specific to flagellar regulation; rather they behave more like unregulated, constitutive promoters (Fig. S12, S13). This result suggests that the observed temporal shift in flagellar gene activity may be driven by global intracellular factors rather than flagellum-specific ones. Indeed, the signature observed in the correlation between elongation rate and Class-2 activity (see Fig. S13) is consistent with the scenario described by Kiviet and Nghe et al. [38], in which the dominant noise source fluctuations are global cellular processes.

Recent work has observed heterogeneity in ribosome density across populations of *E. coli* cells [39–42]. While Pavlou et al. found no immediate association between ribosome abundance and unequal division [41], several super-resolution studies suggest that ribosome heterogeneity arises from asymmetric partitioning during cell division [39, 40, 43]. They demonstrated such asymmetry stems from large clusters of ribosomes that are asymmetrically distributed at cell division, leading to differences in growth after division. Such a pronounced effect would not be possible if each ribosome was independently partitioned according to a binomial distribution, which, due to their large number, would result in a highly symmetrical distribution between daughter cells [15, 44]. With these results in mind and given that the ribosome synthesis is an upstream process relative to flagellar gene expression, we proceeded to analyze the symmetry between cells following cell division as well as the heritability of flagellar gene activity and growth rate fluctuations from one generation to the next.

First, we asked whether after each division the two new daughter cells differ systematically in growth and flagellar gene activity. We compared the daughter with the new pole and the daughter that inherited the old pole (cell adjacent to the closed end of the channel inherits the old pole) (Fig. 3A). We found a strong bias: the daughter with new pole has a greater elongation rate than the daughter with the old pole. This bias is even stronger when the daughter with the new pole also has a greater promoter activity than its sister (Fig. S14).

**Fig. 3.**
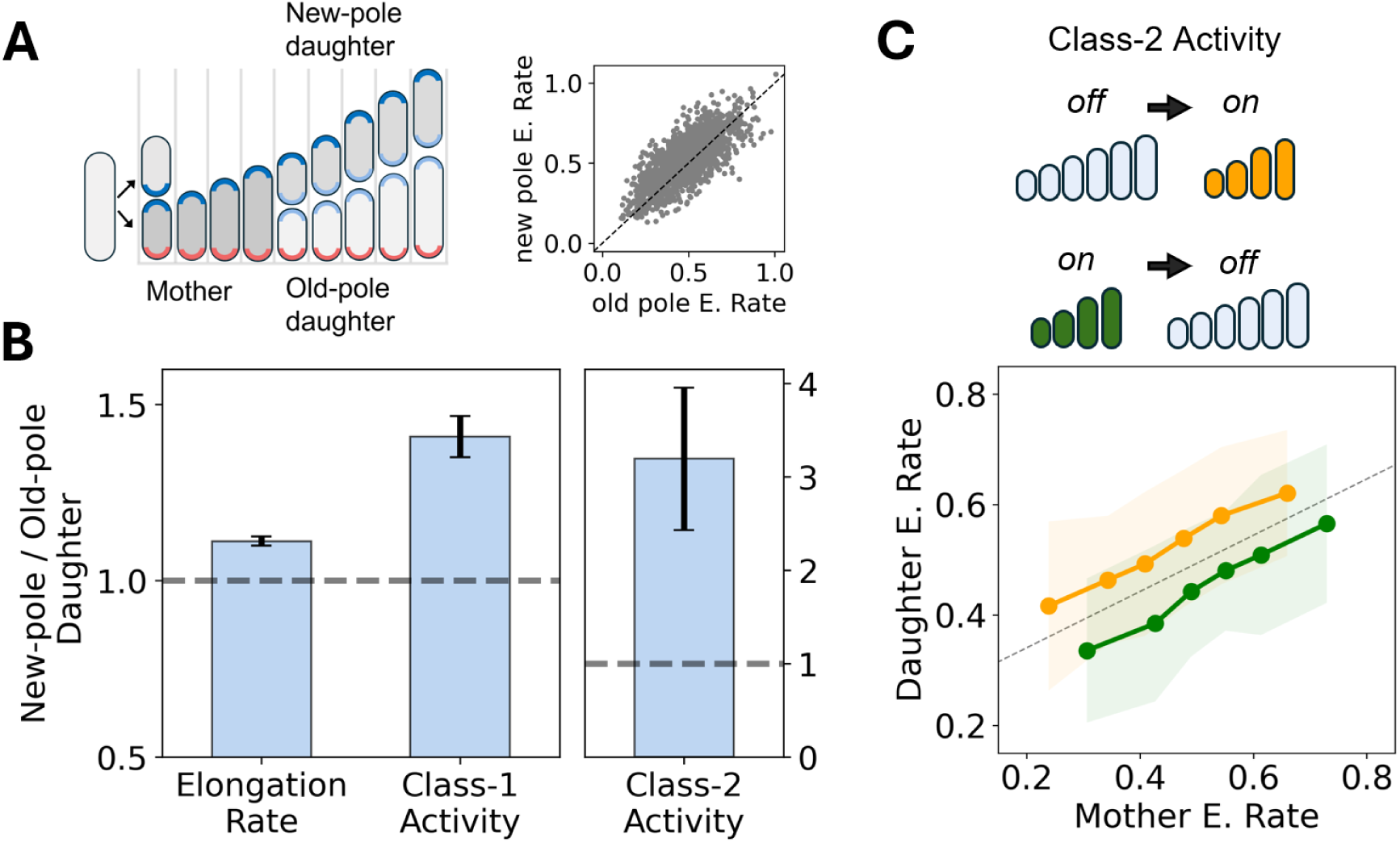
Short-timescale shifts in Class-2 activity correlate with immediate changes in growth. (**A**) Left: Schematic of a dividing mother cell producing two daughters, one inheriting the new pole (red pole) and the other retaining the old pole (blue pole). Scatter plot of elongation rate in sister cells (new-pole / old-pole daughters. (**B**) Bars show the mean ratio of new-pole to old-pole daughters for elongation rate, Class-1 activity, and Class-2 activity. The gray dotted line indicates a ratio of 1, corresponding to equal values between new- and old-pole daughters and is also the mean ratio obtained from a null model based on label-shuffling permutations between the daughters. Error bars indicate 95% confidence intervals. N = 1669 pairs of the WT flagellar expression strain. (**C**) Elongation rates across two consecutive generations for cells experiencing an abrupt change in Class-2 activity: *off* to *on* (yellow) and *on* to *off* (green). For both cases, we plot the binned *ER* of the daughter cell (generation g + 1) against the *ER* of the previous generation (g). A positive change in activity (*off* to *on*) is accompanied by an increase in *ER* across generations compared to the dotted line, reflecting faster cell growth in generation g+1. A decrease in activity (*on* to *off*) is associated with a reduced *ER* compared to the dotted line in generation g+1 (*ER* is lower at g+1 than at g). The number of divisions for each subset is ∼300. The shaded regions correspond to the standard deviation for each bin, and the dotted line shows the linear fit for the entire dataset, irrespective of activity, representing a null model for *ER* across divisions.

To quantify this bias, we computed, at each division, the ratio of elongation rate and Class-1 and Class-2 promoter activity between new-pole and old-pole daughters. All ratios are clearly above 1 (Fig. 3A), showing that new-pole daughters tend to grow faster and exhibit higher flagellar gene activity than old-pole daughters. This observed bias is consistent with previous work showing that old-pole regions lose gene expression activity [45], and that aging at the old pole enhances phenotypic heterogeneity [46]. More importantly it agrees with observations that ribosomes are denser in daughters with new poles, where higher ribosome levels are associated with higher elongation rates [40]. Together, these observations support the hypothesis that differences in growth rate between sister cells are governed by difference of growth factor distribution at division.

We then focused exclusively on cells that exhibited changes in flagellar gene activity across division (from mother to daughter) and measured the corresponding changes in elongation rate. We first selected the cells that transitioned from *off* to *on* activity across division (Fig. 3B, upper right) and compared their elongation rates (Fig. 3B, yellow). Our results indicate that the increase in activity across divisions is associated with an increase in the elongation rate that is greater than for cells with no transition (*off* - *off*, dotted line). By contrast, cells showing a transition from *on* to *off* activity (Fig. 3B, green) fall below the dotted line.

Together, these observations show that cell division systematically generates asymmetry in both growth and flagellar gene activity and that transitions in flagellar activity across division are accompanied by concordant changes in growth rate, indicating that both traits are modulated by short-timescale global fluctuations.

Next, we asked whether these observations could be strain dependent. The MC4100 E. coli strain is known to exhibit long-timescale oscillatory dynamics in cell size, distinct from MG1655 [44]. We found that this slow, ∼10-generation oscillation is present in MC4100’s cell size but is absent from its elongation-rate autocorrelation (Fig. S15A, B). Instantaneous elongation rate remained positively correlated with Class-1 promoter activity in MC4100, comparable in magnitude to MG1655, indicating that the short-timescale growth-expression coupling itself is not strain-specific. However, this correlation was markedly attenuated at the level of Class-2 flagellar activity in MC4100 specifically (Fig. S15C), suggesting that the discrepancy arises in the transmission of growth-coupled fluctuations through the flagellar regulatory cascade rather than in the growth-coupling mechanism itself.

Next, we explored whether the single-cell behavior observed in Fig. 1, namely the violation of the population-level growth law, is specific to the flagellum system or if it constitutes a more general phenomenon observable at the single-cell level. To this end, we repeated our experiments using unregulated synthetic expression systems that govern the expression of the yellow fluorescent protein, Venus, by means of synthetic constitutive promoters with various strengths. Although our bulk measurements (Fig. 4A, diamonds, Fig. S16C) as in [5] experimentally confirm the standard growth law, where growth rate decreases as unnecessary protein expression increases, binning analyses of instantaneous growth rate measurements revealed a strikingly different behavior at the single-cell level. Using a validated model relating growth-rate reduction to unnecessary-protein burden, based on the ribosome-allocation theory of Scott et al. [6] and calibrated for GFP-family proteins by Bienick et al. [47], we estimate that mVenus constitutes approximately 0.6–4.0% of dry cell weight across our promoter series (Table S3). As with the flagellum system, our analysis shows that an instantaneous increase (decrease) in fluorescent protein synthesis is accompanied by a faster (slower) elongation rate (Fig. 4A; Fig. S16A, B; cross-correlation between elongation rate and activity in Fig. S17). The result in Fig. 4A appears to contradict the growth law, which posits that the increased synthesis of unnecessary proteins, such as the fluorescent proteins in our experiments, depletes ribosomal availability and should therefore reduce growth rate. However, our single-cell analyses reveal an opposite behavior, where transient boosts of growth accompany increases in protein synthesis. Consequently, we hypothesized that the observed behavior likely arises from an out-of-balance growth process over short timescales during which a surplus (or deficit) of beneficial growth factors is redistributed at division from the mother at no cost to the daughter cell. The process is out-of-balance because the post-division abundance of growth factors (such as ribosomes) reflects unequal inheritance rather than balanced synthesis.

**Fig. 4.**
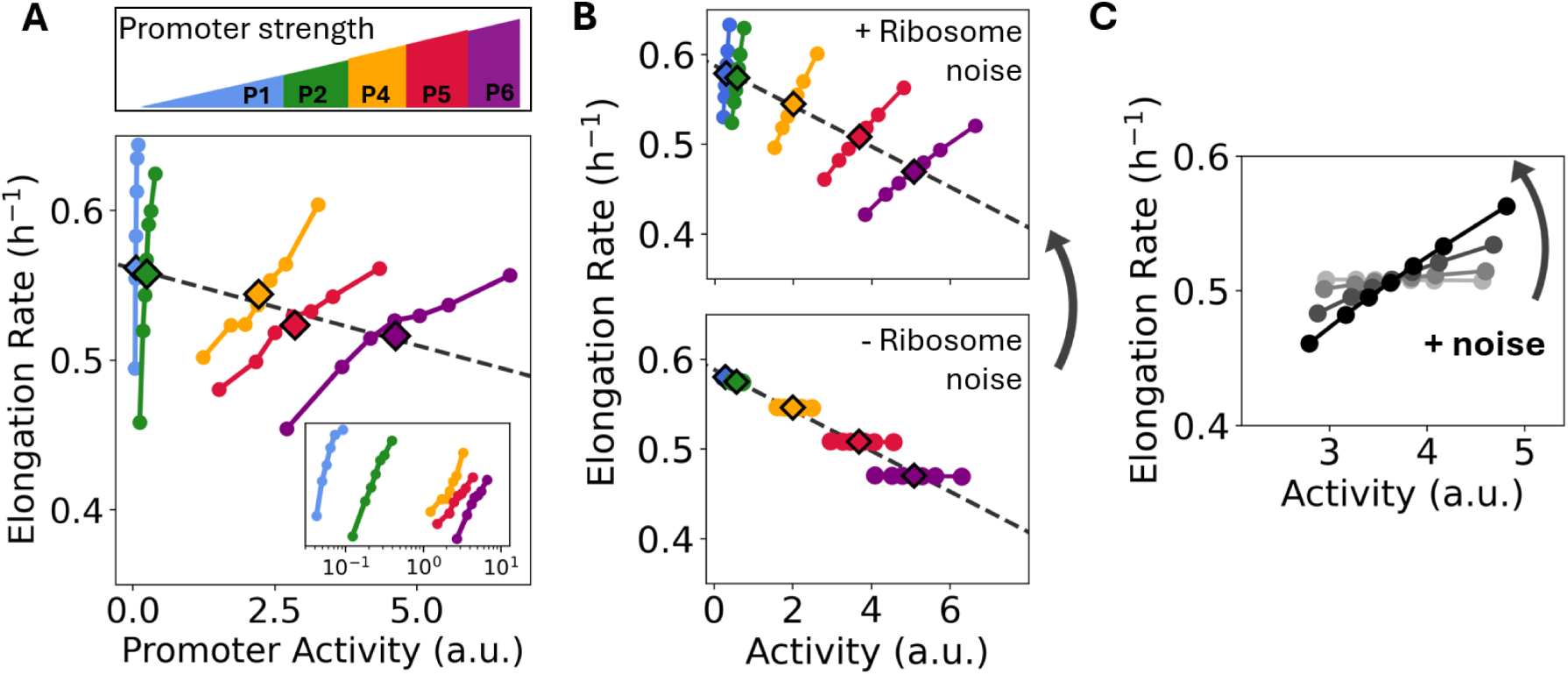
Relationship between instantaneous growth and promoter activity in single cells versus population for different levels of constitutive expression of unnecessary proteins. (**A**) For a set of strains expressing the fluorescent protein Venus, driven by promoters of varying strengths (P1– P6, as indicated at the top), we calculated the mean elongation rate as a function of promoter activity. The diamond markers represent the average for the entire population of each strain, and the black dotted line indicates the linear fit through these population averages. The colored dots show the binned single-cell data for each strain. In each strain, cells exhibit a positive correlation between elongation rate and constitutive promoter activity. The inset shows the same data with the x-axis plotted on a log scale. (**B**) (top) Single-cell simulations for increasing average allocation fraction to unnecessary protein expression 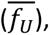 capturing both the decreased elongation rate with increasing activity at the population level across conditions (diamonds), as well as the binned data, that shows the increase in elongation rate with activity at the single-cell level within conditions (colored dots). (bottom) Single-cell simulations of elongation rate as a function of activity with identical parameters to (top), except without noise due to unequal partitioning of ribosomes at division (*σ_R_*=0), showing that the positive slope between elongation rate and activity seen at the single-cell level (top) is driven by fluctuations in ribosome concentration at the single-cell level. (**C**) Simulations were performed for a fixed fraction of unnecessary proteins, *f_U_*= 0.073, and varying noise amplitudes of ribosomes due to unequal partitioning *σ_R_*=(0,0.02,0.04,0.06) (light gray to black). For each condition, the resulting scattered data of activity versus elongation rate was binned as in the experiments. All simulations were implemented using rules for proteome allocation and division timing from [14], and other simulation parameters are described in SI Section 1.

To test whether this coupling extends beyond our own strains and experimental conditions, we reanalyzed publicly available single-cell data from two independent studies of *E. coli* constitutively expressing a fluorescent protein: Wang et al. [33], in MG1655 and B/r, and Tanouchi et al. [44, 48], in MC4100 at 25°C and 37°C. In all cases, we found the same positive, short-timescale correlation between instantaneous elongation rate and expression (Fig. S18) indicating that the out-of-balance growth process we describe is not limited to our strains or growth conditions, but is a broader feature of single-cell *E. coli* physiology.

To evaluate whether the proposed out-of-balance growth mechanism can account for these observations, we performed single-cell simulations using a computational model (Fig. 4B) that incorporates all the key elements that govern the growth laws as commonly defined from population measurements (Simulation details in the Supplementary Materials). Importantly, the model also includes the stochastic redistribution of growth factors such as ribosomes and possibly other factors from the mother to daughter cells at division. We find that our binning procedure shows the same positively correlated relationship between growth rate and synthesis of unnecessary proteins observed experimentally (Fig. 4B, top). Moreover, the slope of this relationship becomes steeper as the noise in the distribution of growth factors becomes larger (Fig. 4C). In the absence of noise, the relationship shows no correlation between growth rate and protein synthesis (Fig. 4B, bottom). In contrast, this correlation exhibits increased steepness with higher noise levels, indicating that stochastic partitioning of growth factors amplifies transient out-of-balance states.

This effect shows that instantaneous growth and expression changes are mainly dominated by inheritance noise rather than by the steady-state burden predicted by the growth law. In our model, single-cell growth rate is positively correlated with protein synthesis, so any mechanism generating molecular growth factors heterogeneity across generations will give rise to Simpson’s paradox when comparing single-cell and population-level trends.

Altogether, these simulations illustrate that both behaviors can coexist: the growth burden caused by the synthesis of unnecessary proteins as seen at the population level, and the out-of-balance growth observed at single-cell level, which mediates instantaneous, burden-free synthesis of unnecessary proteins.

Rather than identifying the precise molecular source of heterogeneity, our goal is to examine the consequences of fluctuations of unspecified growth factors for balanced growth models. We therefore propose that these fluctuations underlie an intergenerational mechanism for cost-free (“pre-paid”) protein production. We argue that mother cells can pass molecular growth factors to their daughters, either through excess of their production or asymmetric division, thereby giving these daughters an advantage in synthesizing costly proteins such as the flagellum.

## Discussion

Kim et al. [20] found that flagellar genes in E. coli are activated in stochastic pulses. Subsequently, some of us proposed that these pulses are generated by means of an inhibitor of the Class-1 transcription factor, called RflP (formerly YdiV), which creates an ultrasensitive switch that acts as a digital filter to remove small-amplitude input temporal fluctuations in Class-1 expression [24]. While this molecular mechanism demonstrates that pulses can arise without feedback, it does not address how such pulses influence cellular growth. This switch may also contribute to strain-specific differences: we found that in MC4100, the short-timescale coupling between elongation rate and expression, intact at the Class-1 level and in constitutively expressed proteins across strains (Fig. S15C, Fig. S18), is markedly attenuated at Class-2. Rather than reflecting a general difference in growth-factor partitioning, which we show is comparable to MG1655 at the Class-1 level, this result suggests that the flagellar pulse-generating circuit itself may differ across strains in how, or whether, it exploits this generic growth-coupling noise.

Knowing that these pulses are not an essential feature of flagellar biosynthesis, especially since some strains can be genetically modified to produce flagella steadily [49], this raises the question of the biological advantage of such a pulsing dynamics in wild-type strains. Here, we address this question through time series analyses to demonstrate that pulses of flagellar gene activity accompany sudden growth boosts that transiently alleviate the growth burden associated with flagellar gene expression. By temporarily mitigating this burden, we speculate that some single cells may more effectively explore and colonize new environments as the cell does not have to immediately ‘pay’ for the cost of flagellum expression. In particular, it would be especially interesting to investigate whether this advantage is more pronounced in nutrient-depleted environments where seeder bacteria with a faster growth rate would also be the ones producing flagella. The fact that the relationship between growth rate and flagella expression, or more generally fluorescent protein expression, at the single-cell level contradicts the population-level trend across strains observed in bulk measurements (Fig. 1, Fig. 4) is reminiscent of Simpson’s paradox and arises from the separation of timescales inherent to these distinct analyses of the same sample [17, 18, 22]. Similar time series analyses based on the separation of timescales may also reveal that out-of-balance growth processes serve as a general mechanism for temporarily mitigating the cost of expressing other energetically demanding endogenous genes [21].

## Acknowledgments

We thank Maximino Aldana, Mark Kim and Enrique Balleza for early discussions, advice and guidance, the Cluzel lab members, Fotios Avgidis for critical reading of the manuscript, and Camilla U. Rang for drawing our attention to the importance of the old versus new pole bias. AI-assisted technologies were used to prepare the manuscript. Funding: This work was performed, in part, at the Center for Nanoscale Systems, a member of the National Nanotechnology Infrastructure Network, which is supported by the National Science Foundation under award no. ECS-0335765. The Center for Nanoscale Systems is part of Harvard University. This work was supported in part by grant 1615487 from the National Science Foundation and by the United States Government, ARPA-H. The views and conclusions contained in this document are those of the authors and should not be interpreted as representing the official policies, either expressed or implied, of the United States Government.

## Materials and Methods

### Strains

For a complete description of the strains see, Table 2 in the Supplementary Materials. Briefly: MG1655 flagella reporter strains (E98KFD and E98KFD-Pro) carry the gene for yellow fluorescent protein (mVenus NB) inserted downstream of the Class-1 native promoter of the flhDC operon. At the attB site, a Class-2 promoter, fliFp, controls the activity of the cyan fluorescent protein SCFP3A. The E98KFD-Pro strains differ from E98KFD only in that the native Class-1 promoter is replaced with constitutive synthetic promoters with various strengths (P1: Pro2, P2: Pro4, P3: Pro5, from [37]). The strain construction is described in detail in [20].

Strains with constitutive expression of fluorescent proteins (MGPro-Venus; [50]): each strain carries a low-copy plasmid with an SC101 origin [51], in which a synthetic promoter [37] controls the expression of the sequence of the mVenus NB yellow fluorescent protein [52]. We renamed the promoters according to their reported strength and measured fluorescence level (P1: Pro2, P2: Pro4, P4: ProB, P5: ProC, P6: ProD).

MC*: The classic strain MC4100 [44, 53] was modified to correct for its flh5301 point mutation by replacing flhD with the same gene sequence as in MG1655 in order to recover its flagellar expression. We also removed the insertion element IS5 upstream of flhDp so as to keep Class-1 levels consistent with those in MG1655. We introduced the mutation (MotA E98K) to make the strain non-motile. Finally, the strain carries the Class-1 and Class-2 fluorescent reporters as in the MG1655 E98KFD strain. To implement mutations, we used a scarless chromosomal engineering strategy described in [20], and all transformations were performed via electroporation.

### Growth Media

Microfluidic experiments were conducted using a modified version of Teknova’s MOPS-EZ Medium: MOPS and Supplement EZ at 1x concentration; 0.4% glycerol serving as the carbon source, phosphate at 1.32 mM. Additionally, we added surfactant (Pluronic F-108, Sigma-Aldrich) at a final concentration of 0.85 g/L. Overnight cultures were prepared using LB medium.

### Microfluidic device

We used a Mother Machine device customized for the lab setup. The master mold chip has 4 independent main feed channels where growth media flows, feeding smaller microchannels with a closed end (length × width × height = 25 μm × 1.2 μm × 1.2 μm).

We fabricated the microfluidic devices by casting a 10:1 mixture of degassed PDMS and curing agent (Sylgard 184, Dow Corning) onto the mold, curing it overnight at 65 °C. The individual chips were then removed from the mold, and inlets and outlets were created using a 0.75 mm biopsy punch, then they were bonded to a glass slide (25 mm × 40 mm No. 1.5 coverglass (VWR)) using a plasma cleaner and kept at 65 °C for another hour.

### Experimental Setup

Individual colonies were cultured overnight at 30°C, then concentrated by centrifugation for 1 minute at 8000 g and loaded into the device by pipetting. After loading the device, the sample was left for 1 hour at 30°C to increase the likelihood of the cells entering the channels by diffusion. Then the trapped cells in the channels began to grow exponentially. We flowed the media through the microfluidic chamber using Tygon tubing (VWR; inside diameter 0.02 inch × outside diameter 0.06 inch), and the flow-through was collected from an outlet via a second tubing into a waste beaker. Growth medium was first pumped at a rate of 20 µl/min for at least 1 hour to allow the inlets and outlets to be cleared.

Subsequently, the flow rate was decreased to 9 µl/min, and the temperature was maintained at 30°C. We allowed approximately 3 hours for the cells to adapt before initiating the measurements. The time-lapse images were acquired every 5 minutes on a Zeiss Axiovert 200M microscope equipped with a Plan-Apochromat 40x/1.3 Oil Ph3 Objective, a CCD camera (Hamamatsu C4742-98-24ERG) and a fluorescence excitation LED illumination source (SOLA SE II, Lumencor). We collected images from the phase-contrast and epifluorescent channels (yellow, blue, and/or red protein filters, depending on the strains). The typical duration of the experiment was 30-60 hours.

### Microscopy Image Processing

We used the software BACMMAN (BACteria in Mother Machine Analyzer) [54] for cell segmentation and tracking. Using this tool, we customized various parameters to process images according to each specific strain and condition, and then manually corrected any segmentation errors.

### Elongation rate and promoter activity calculations

To obtain the elongation rate *ER*(*t*), we determined the first derivative of individual cell sizes *S*(*t*) (from birth to division) using a Savitzky–Golay filter (from Python’s SciPy library) with a window of 7 data points (35 minutes). This derivative was then normalized by dividing by the cell size, resulting in the expression *ER*(*t*) = 1/*S*(*t*) × *dS*(*t*)/*dt*.

As a proxy for the promoter activity, we used *PA*(*t*) = 1/*S*(*t*) *dF*(*t*)/*dt*, where *S*(*t*) is the cell size (number of pixels at time t), and *F*(*t*) is the total fluorescence. As in previous literature [17, 20, 55–57], we considered the rate normalized by cell size, thereby adjusting the production rate so that it is expressed per chromosomal equivalent. By normalizing in this way, we focus on the gene’s specific activity, rather than the cell’s total output.

Given that the total fluorescence *F*(*t*) is the mean intensity of the cell, *C*(*t*), multiplied by its size *S*(*t*), one can show that *PA* can also be written as *PA*(*t*) = *C*(*t*) × *ER*(*t*) + *dC*(*t*)/*dt*. We used this equation to calculate the *PA*, as the mean fluorescence effectively represents the intracellular concentration of the fluorescent signal, which changes more continuously and is less affected by the sudden halving associated with cell division, making the calculation of the derivative less prone to artifacts. We smoothed *C*(*t*) with a window of 35 minutes, which we slid over the time series with an increment of 5 min. We calculated the time derivative using the Savitzky–Golay filter, also with a window of 35 minutes.

### Binning

To analyze the single-cell relationship between *ER*(*t*) and *Activity(t)* across all acquired time points, we first visualized the data using a scatter plot. We then binned the data based on *Activity*(*t*) values, as the series displayed a clear pulsatile behavior and distinct mean values across strains. Each bin contained an equal number of data points. Finally, for each bin, we calculated the mean values of both *ER*(*t*) and *Activity(t)*.

### Pulse-to-pulse analysis

We defined Class-2 pulse-to-pulse intervals by empirically choosing a threshold to determine the on (above the threshold, the pulses) and *off* states (below the threshold). Then, for each sequence of on-off-on states, we identified the two maxima within the Class-2 activity time series and included the 15 timepoints before and after the maxima for each time interval.

We grouped those intervals so that they had similar durations and rescaled them from 0 to 1. Following the rescaling, we applied a sliding window to compute the average of the Class-2 activity within these normalized intervals, which allowed us to move systematically over the data points. This method provided us with a smoothed representation of the Class-2 activity, highlighting the significant changes. To examine the ‘instantaneous’ relationship between the pulses and other quantities, we analyzed the same time intervals within the same cells initially defined by the Class-2 activity for the elongation rate, division time, etc. We then rescaled the intervals and used the same window size as for Class-2 activity to average them together. The averages of these quantities for different pulse-to-pulse duration intervals are presented in Fig. S6.

### Autocorrelation and Cross-Correlation Calculations

To quantify the cross-correlation between elongation rate *ER*(*t*) and flagellar activity *A*(*t*), we first subtracted the mean from each time series *ER*^′^(*t*) = *ER*(*t*) − ⟨*ER*⟩, *A*^′^(*t*) = *A*(*t*) − ⟨*A*⟩, and then used the function ‘correlate’ from the SciPy library in Python. The normalized cross-correlation is defined as

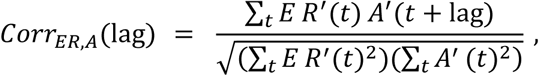

where *l* is the time lag. In practice, we evaluated the numerator using ‘correlate’ function from SciPy library in Python and normalized by the square root of the product of the zero-lag autocorrelations, by using the same function.

With this convention, the order of the arguments is important: in *Corr_ER_*_,*A*_(l), a maximum at negative *l* indicates that changes in *ER* tend to precede changes in *A*, whereas a maximum at positive *l* indicates that activity tends to precede elongation rate.

To average the cross-correlation for different lineages from the same strain, we adjusted the lag by dividing it by the average division time of the series. This rescaling ensures that fluctuations due to cell division are aligned across the different series. We applied the same methodology for the autocorrelation function.

## Author Contributions

P.C. and M.G.-A. conceived the study and analyzed the data. M.G.-A. performed experiments and statistical analyses; J.C.K. performed the simulations. P.C. supervised the project. M.G.-A. J.C.K and P.C. wrote and prepared the manuscript.

## Supplemental Materials

## Experimental validation of functional flagella expression in reporter strains

We performed a motility plate assay to qualitatively verify that the strains we claim have different levels of flagellar expression display phenotypes consistent with their promoter strengths as reported by Davis et al. [37]. On low-agar plates, we inoculated strains with the native Class-1 promoter was replaced by Pro2, Pro4, or Pro5 (P1-P3 in the main text), together with the following controls: the wild type MG1655, MotA E98K MG1655 (strain with a mutation that disables the rotation of the flagella [35, 36]), MotA E98K Pro4, Pro4 ΔfliC, and MG1655 strain carrying the IS5 insertion (a motile strain known to have elevated flagellar expression [58]). One microliter of liquid culture was spotted onto the plate and incubated for ∼12 h at 30 °C. Motile strains formed characteristic swarming rings (Fig. S3A). The strains with weaker promoters produced smaller swarm diameters than those with stronger promoters, in the same order expected from Davis et al. and consistent with our fluorescence measurements after introduction of the reporter. There was no growth ring for the strains carrying the MotA E98K nor ΔfliC mutations. This confirms that the reporter activity tracks the level of functional flagellar expression across strains.

Then, to address whether cells are flagellated and motile in the mother-machine channels, we compared the wild type MG1655 and MotA E98K MG1655 under identical microfluidic conditions. Cells were loaded in stationary phase and allowed to recover before time-lapse imaging. In the wild-type strain, we observed that ∼35% of lineages left the growth channels during the experiment after 20 hrs, whereas non-motile cells did not leave the channels. Because the only difference between these strains is the MotA E98K mutation, this strong differential escape behavior indicates that cells in the channels do assemble flagellar filaments and motors, and when motors are activated, the resulting motility is sufficient for cells to swim out of the channels. In Fig. S3B we show one channel of the mother machine filled with cells outlined in blue, which suddenly leave the channels. Together with the motility plate assay, this supports that our reporter signal corresponds to expression of functional flagella under the conditions of our experiments.

## Simulation details: stochastic proteome allocation model with division-coupled noise

Single-cell simulations were implemented using rules for proteome allocation and division timing from ref. [14], allowing for analysis of instantaneous proteome allocation and growth rate at the level of the individual cell. For completeness, here we include a description of all relevant equations required to define the model, but for a detailed derivation and model justification, we refer readers to ref. [14]. Each individual cell can be described by a set of state variables which include its coarse-grained proteome allocation to five sectors of proteins grouped together based on function and defined by their mass fractions *φ_i_* for *i* ∈ {*R*, *P*, *X*, *Q*, *U*}, specifically ribosomal *(R*), metabolic (*P*), division (*X*), housekeeping (*Q*), and exogenously expressed “unnecessary” (*<u>U</u>*) proteins. These sectors are coupled to the amino acid mass fraction of the cell, *a*. The ribosomal (*ϕ_R_*) and amino acid (*a*) mass fractions together set the exponential growth rate, *κ* = d ln *V* /d*t*, given by:

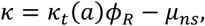

where *μ_ns_* is the non-specific degradation rate constant and 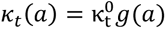 is the translational efficiency, which is dependent on *a* such that translation becomes significantly attenuated at low amino acid levels (specifically *g*(*a*) = (*a*/*a_t_*)^2^/[1 + (*a*/*a_t_*)^2^]). The dynamics of each sector are given by:

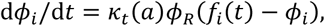

where *f_i_*(*t*) denotes the fraction of the total translational flux devoted to sector *i*. Both *f_R_* (*t*) and *f_X_*(*t*) obtain their time dependence through their dependence on other dynamic quantities, specifically *a*(*t*) and 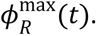 These expressions are given by:

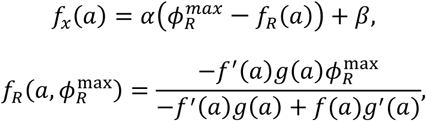

where *f*(*a*) = 1/[1 + (*a*/*a_n_*)^2^], 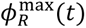 is a dynamic parameter constraining the maximum allocation to ribosomal proteins, and here *y*^′^(*a*) denotes the derivative of *y* with respect to *a*. *f_Q_* is taken to be invariant to nutrient changes, thus imposing the additional constraint 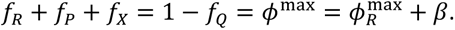 The dynamics of *f_U_* are discussed below. The dynamics of *φ_i_* are coupled to the dynamics of *a*, which are given by:

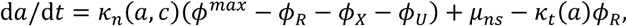

where 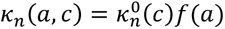 is the nutritional efficiency and *c* is the extracellular nutrient quality. Finally, a division event occurs when a sufficient number of *X* proteins have accumulated. The dynamics of *X* are given by:

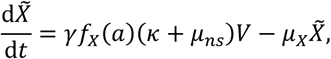

where here we track the normalized quantity *X̃* = *X*/*X*_0_such that a cell divides when *X̃* = 1, *μ_X_* denotes the division protein degradation rate, and *γ* is a conversion factor relating protein mass fraction to protein number (see ref. [14] for details). To match experiments, at each division event a single daughter cell was simulated which inherited the intracellular composition of the mother cell, maintaining the overall population size.

Previous experimental work has shown that the major driver of heterogeneity in ribosome concentration across single cells is not stochastic production, but rather unequal partitioning of ribosomes at division due to the formation of larger clusters [39]. Motivated by this, we modeled the redistribution of ribosomes at division through a stochastic differential equation in which perturbations to the maximum allocation to ribosomes only occurred at division times, *t_d_*, then decayed slowly with a mean-reversion timescale *τ_R_* ≫ 0. Specifically:

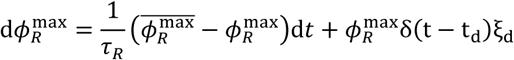

where 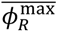 is the mean maximum allocation to ribosomes, *δ*(·) is the Dirac delta function, and 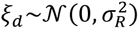 denotes random jump magnitudes drawn from a Gaussian distribution of mean 0 and variance 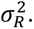 In the absence of clustering, *σ_R_* = 0.

In contrast, we assumed the main driver of heterogeneity in unnecessary protein allocation is due to stochastic gene expression. As such, we modeled noise in unnecessary protein expression by assuming it continuously fluctuated around its mean allocation value 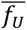 with timescale *τ_U_* and with volatility coefficient *σ_U_*, yielding:

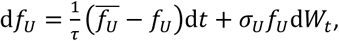

where *W_t_* denotes a Wiener process. Assuming constant cellular density, protein mass fraction is proportional to protein concentration, allowing us to rewrite our previous definition of promoter activity as: 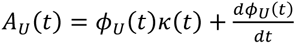 where *ϕ_dt_*(*t*) is the mass fraction of unnecessary proteins and *κ*(*t*) is the single-cell growth rate at time *t*. Using this expression, along with our definition of sector dynamics from above 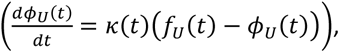 the activity of unnecessary protein expression could then be calculated as:

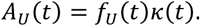

To simulate, we numerically integrated the set of coupled stochastic differential equations detailed above for each cell. The code used to generate all simulation results are available at: https://github.com/josiah-k/Simpsons-paradox. The single-cell growth rate and corresponding activity for 3000 individual cells was computed at intervals of Δ*t* = 3min for 75 h total.

Parameters

Parameters:
|  |  |
| --- | --- |
| Translational efficiency, $\kappa_t^0$ ($\text{h}^{-1}$): | 5 |
| Nutritional efficiency, $\kappa_n^0$ ($\text{h}^{-1}$): | 1.5 |
| Mean-reversion timescale (ribosomes), $\tau_R$ (h): | 6.25 |
| Mean-reversion timescale (unnecessary proteins), $\tau_U$ (h): | 0.0625 |
| Unnecessary protein expression volatility coefficient $\sigma_U$: | 0.3 |
| Ribosome protein expression volatility coefficient $\sigma_R$: | 0.06 |
| Average unnecessary protein allocation fraction, $\overline{f_U}$: | 0.005, 0.010, 0.037, 0.073, 0.108, 0.144 |
| Division protein degradation rate, $\mu_X$ ($\text{h}^{-1}$): | 0.6 |
| Average maximum flux allocation to ribosomes, $\overline{\phi_R^{\max}}$ [14]: | 0.55 |
| Translation attenuation threshold, $a_t$ [14]: | $10^{-4}$ |
| Feedback inhibition threshold, $a_n$ [14]: | $10^{-3}$ |
| Nonspecific degradation rate, $\mu_{ns}$ ($\text{h}^{-1}$) [14]: | 0 |
| Basal allocation to division protein production, $\gamma\beta$ ($\mu\text{m}^{-3}$) [14]: | 1.1 |
| Dynamic allocation to $X$ protein production, $\gamma\alpha$ ($\mu\text{m}^{-3}$) [14]: | 4.5 |

---

**Fig. S1.**
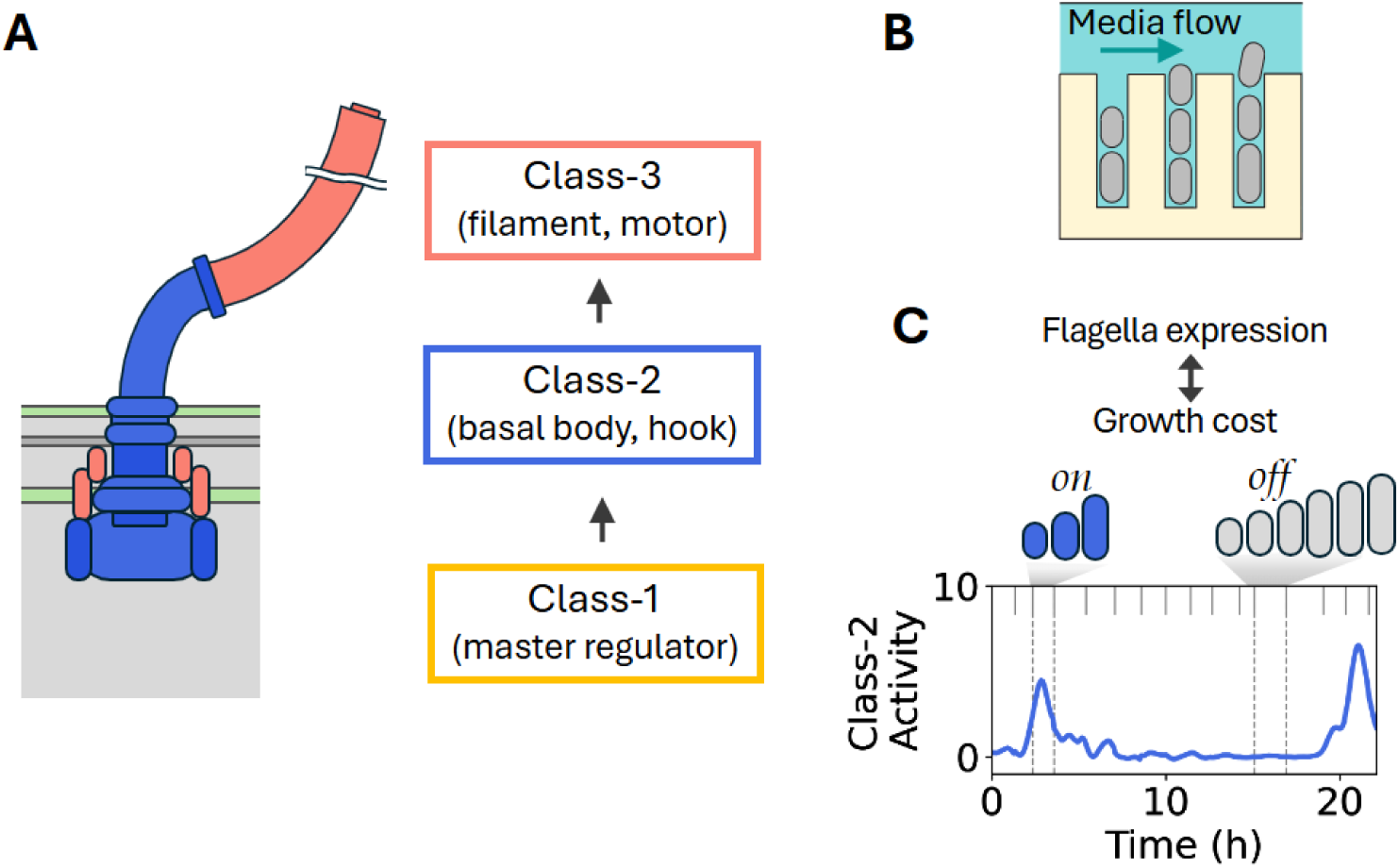
Simultaneous monitoring of growth and pulses of flagellar gene activity in single bacteria. (**A**) Schematic of flagellar components in *E. coli*. A cascade of flagellar promoters coordinates the assembly of the different motor elements. The master regulator (Class-1) regulates the activity of Class-2 promoters that control the genes that encode the basal body and hook molecular elements. A Class-2 gene (*fli*A) drives the transcription of the Class-3 genes, which encode products needed in late flagellar assembly, such as the filament-associated proteins and the motor stator. (**B**) In the Mother Machine device, cells are constrained into channels where they can grow and divide while being fed by a constant flow of fresh medium. The ‘mother’ cell at the bottom of the channel is monitored over many cell divisions using epifluorescence-based time-lapse microscopy. (**C**) Cells growing in the Mother Machine show pulses of Class-2 activity (*on* state) followed by long periods of no activity (*off* state). The activity is calculated from the fluorescence time series controlled by a Class-2 promoter, *fliFp* (described in Materials and Methods; [20]). The vertical ticks show cell divisions; the dashed lines show examples of *on* and *off* states. The cartoon at the top illustrates the different growth behaviors cells exhibit during *on* and *off* states, which yields the relationship between instantaneous flagellar genes activity and growth rate.

**Fig. S2.**
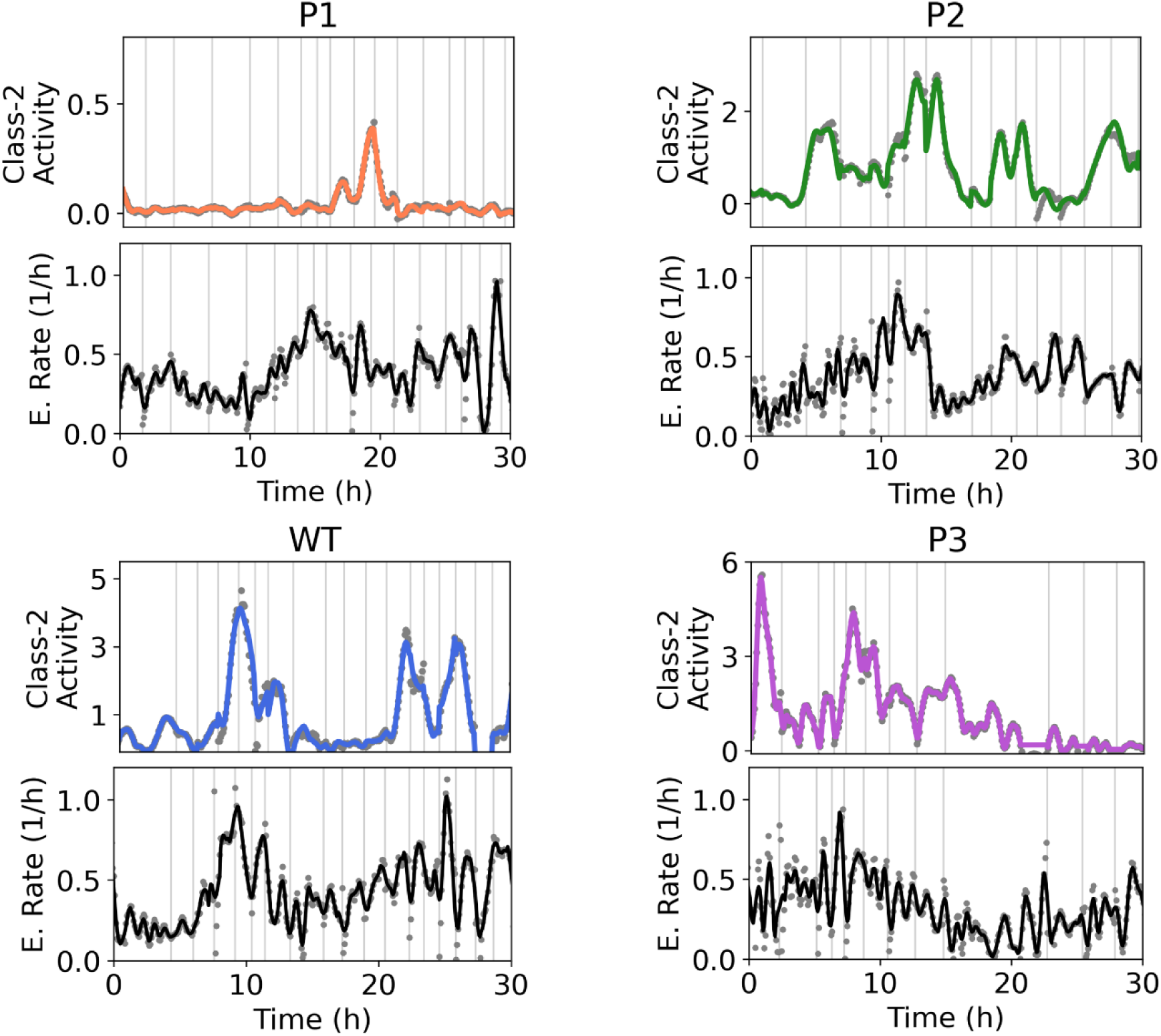
Time series of Class-2 activity and cell elongation rate in flagellar reporter strains. Typical single-cell time series for flagellar reporter strains monitored in the Mother Machine. For each reporter strain (with promoters P1, P2, and P3, which control Class-1 expression) and the wild-type (WT) strain, Class-2 activity and cell elongation rate are shown as a function of time. In each panel, gray dots correspond to the direct calculations of activity and elongation rate, as described in Materials and Methods, and colored lines show the corresponding Savitzky–Golay–smoothed traces (window size w = 35 min). Horizontal lines indicate cell-division events.

**Fig. S3.**
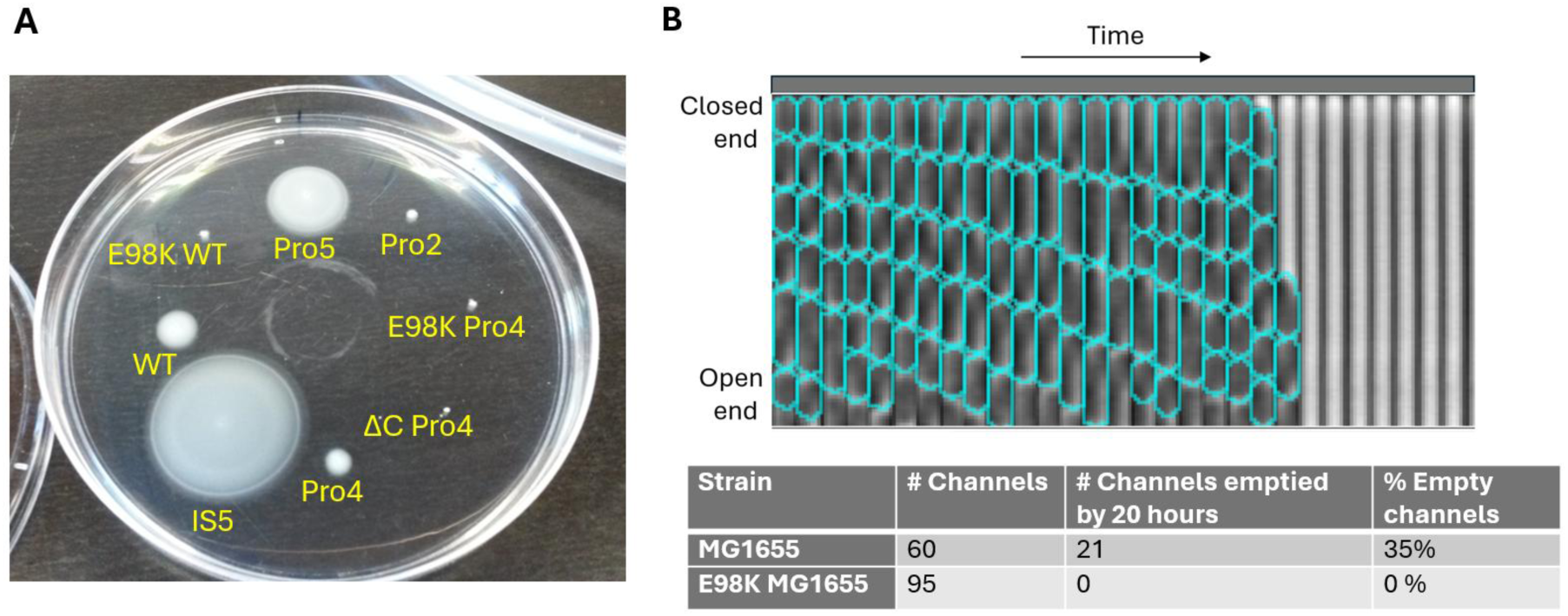
Determining motility of the synthetic flagellar series in soft agar plates and microfluidic device. (**A**) Motility plate assay for strains with synthetic promoters controlling flagellar expression. Each strain (1 µL LB culture) was spotted onto a soft-agar plate (40 mL LB, 0.25% w/v Bacto agar in 150 mm petri dish) and incubated for ∼12 h at 30 °C. Strains in which the native Class-1 promoter was replaced by Pro2, Pro4, or Pro5 (P1–P3 in the main text) were assayed alongside controls: MG1655 WT, MotA E98K MG1655 (non-motile due to impaired motor rotation), MotA E98K Pro4, Pro4 ΔfliC, and MG1655 IS5 (motile strain with elevated flagellar expression [58]). (**B**) Kymograph of a single mother-machine channel showing cells (outlined in blue), imaged every 5 min until they abruptly leave the channel. The table below reports the fraction of lineages that exited the channels; in contrast, the non-motile MotA E98K strain showed no channel escape events. Temperature and growth media were the same of all the other experiments, as described in the Materials and Methods.

**Fig. S4.**
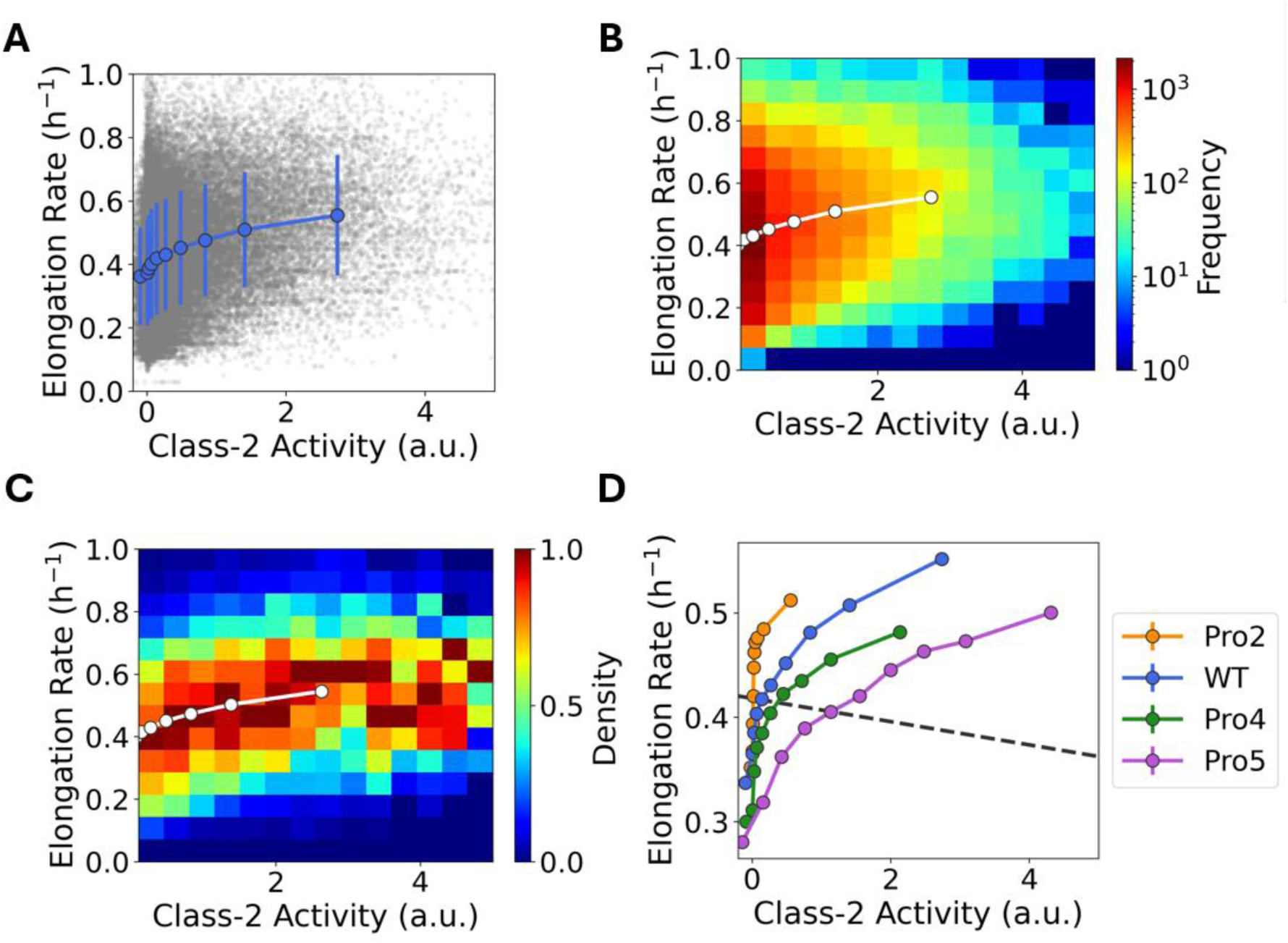
Elongation rate vs Class-2 promoter activity scatter plot and binning. (**A**) The scatter plot in gray illustrates the relationship between elongation rate and Class-2 activity (WT flagellar expression, strain E98KFD), with each point representing a distinct time point. The scatter plot is divided into 10 bins according to the Class-2 activity, each bin containing an equal number of points. The mean and standard deviation for each bin are represented in blue. The dataset consists of a total of n = 57,592 data points, corresponding to n = 107 mother cell lineages monitored over approximately 47 hours. (**B**) The two-dimensional histogram shows the frequency of observations across binned values of Class-2 activity and elongation rate, with the color scale indicating the number of observations per bin (logarithmic scale). (**C**) For each vertical slice (constant activity) from (A), bin counts were normalized between 0 and 1, highlighting the relative density of elongation rate values at each bin of activity. Color intensity indicates normalized density, with 0 corresponding to minimum and 1 to maximum relative frequency within each slice. In (B) and (C) the superimposed points in white represent binned aggregates of the data from (A), illustrating the overall trend. (**D**) Mean binned elongation rate as a function of Class-2 activity, for different strains that have different flagellar expression levels, i.e., strains in which Class-1 activity is controlled by synthetic promoters P1, P2, or P3, and the native promoter, WT. The number of time traces per strain are n = 69, 107, 96, 67 for P1, WT, P2, and P3, respectively, all with a similar duration of ∼47 hours.

**Fig. S5.**
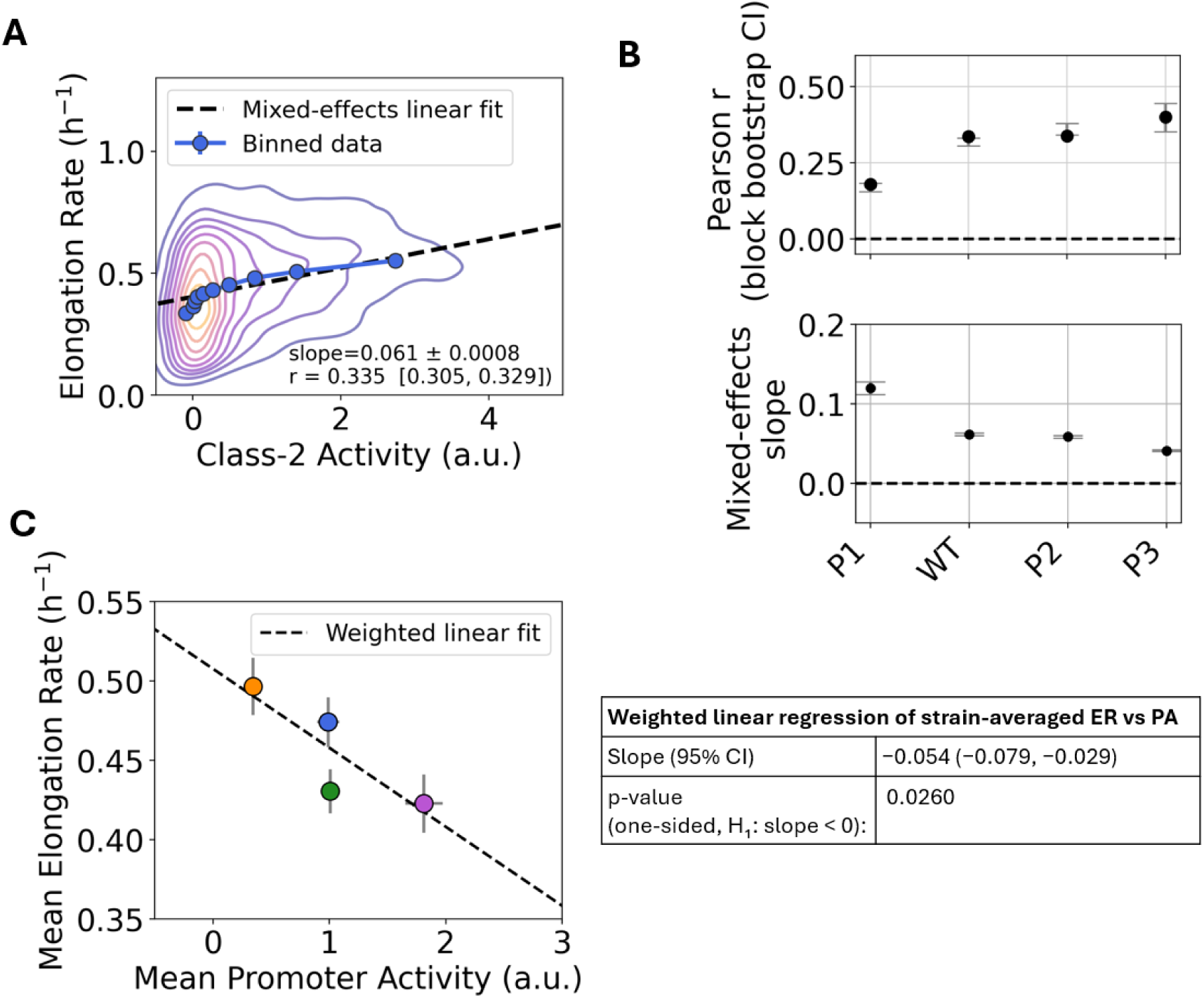
Statistical analysis of single-cell relationships between elongation rate and Class-2 activity. **(A)** The color map shows the density of single-cell measurements of elongation rate versus Class-2 activity, with warmer colors indicating regions containing more data points, for the WT flagellar expression, strain E98KFD. In blue is the previously described binned data. The black dashed line shows the fixed-effect prediction from the mixed-effects linear model (ER ∼ PA with random intercepts per lineage), representing the average single-cell trend after accounting for lineage-to-lineage variability. **(B) Top:** Block-bootstrap estimates of the Pearson correlation between Class-2 activity and elongation rate for flagellar reporter strains (P1, WT, P2, P3), resampling whole lineages to account for within-lineage dependence. Dots show correlations from the original data; vertical lines indicate 95% percentile block-bootstrap confidence intervals. **Bottom**: Mixed-effects linear regression of elongation rate on promoter activity for each strain. Each point shows the fixed-effect slope (change in ER per unit PA); horizontal lines indicate 95% confidence intervals from a mixed model with random intercepts per lineage. The vertical line marks zero slope. (**C**) Mean elongation rate as a function of the mean Class-2 Activity, the error bars show the standard error of the mean and the dotted line shows the weighted linear regression fit (weights proportional to the number of lineages per strain), illustrating the population-level trend across strains. The table summarizes the weighted linear regression of strain-averaged ER on strain-averaged PA, showing the negative slope, its 95% confidence interval, and the one-sided p-value.

**Fig. S6.**
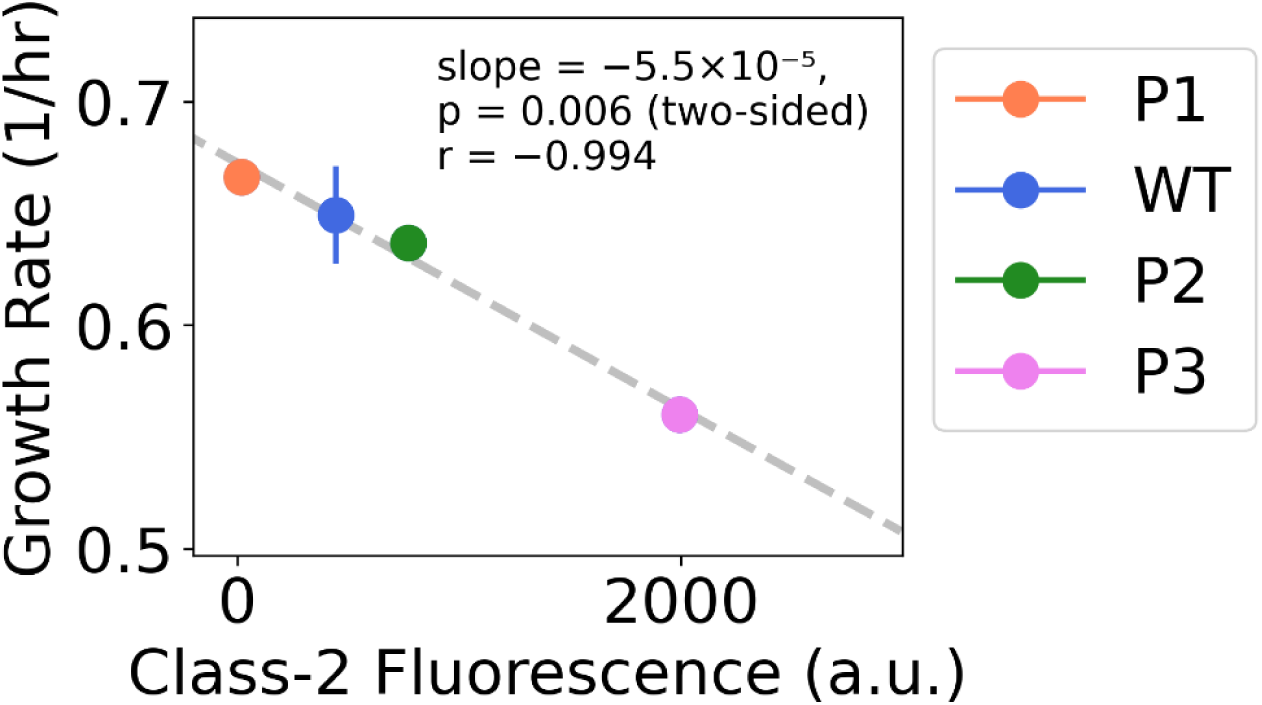
Bulk measurements show a decrease in growth rate when flagellar proteins are overexpressed in *E. coli*. We calculated the growth rate for a set of strains with varying levels of flagellar expression (using different synthetic promoters with various strengths controlling Class-1 expression, P1-P3; see Table S2). Strains were grown in 96-well plates, and their OD600 was measured to calculate the slope of their growth curve during exponential growth. Each strain was also grown in bulk, and flow cytometry was used to measure fluorescence, which indicates the activity of the Class-2 promoter, *fliFp*. The figure shows the growth rate of each strain as a function of the corresponding mean fluorescence. The dashed line represents the best linear fit of the data. The data points reflect the averages for each strain across two biological replicates, and the error bars represent the standard deviation.

**Fig. S7.**
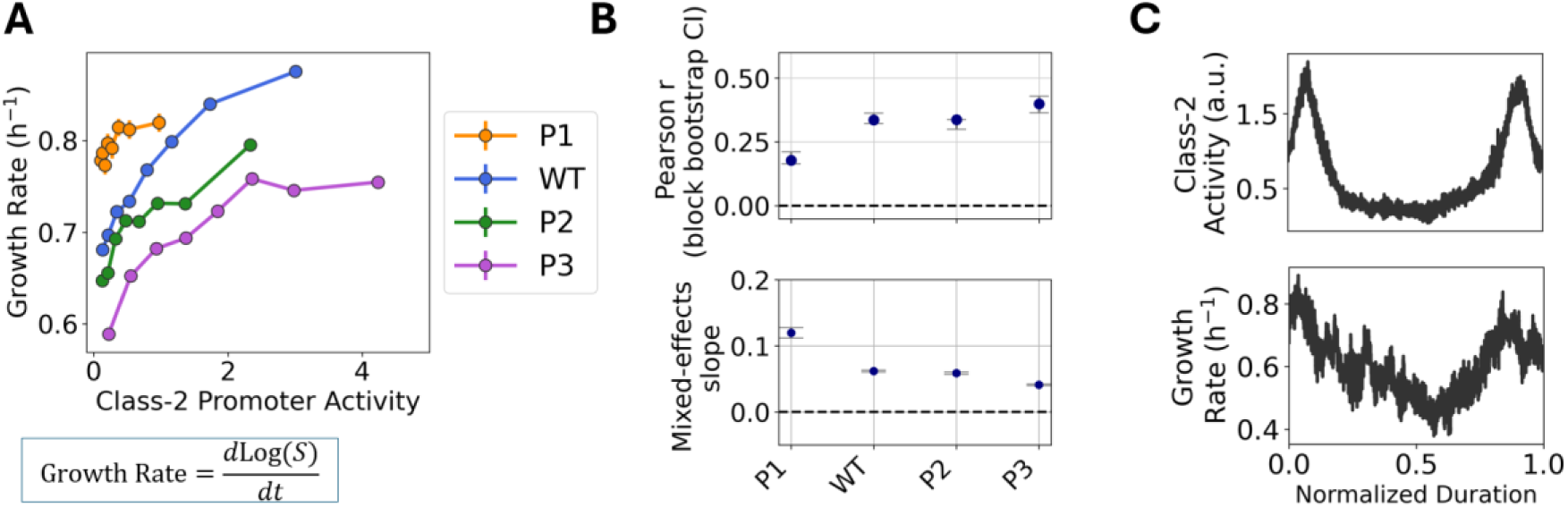
Instantaneous growth rate and flagellar Class-2 promoter activity. (**A**) The mean binned growth rate as a function of Class-2 activity is presented for different strains, each with varying promoters controlling Class-1 activity: synthetic promoters P1, P2, P3, and the native promoter WT. Here, the growth rate was determined as the time derivative of the logarithm of size *S*(*t*), following the methodology of Bakshi et al. [59]. This definition of the growth rate using single-cell measurements was found to be an accurate proxy for the standard bulk growth rate obtained from OD measurements. The number of time traces per strain is 69 for P1, 96 for P2, 107 for WT, and 67 for P3, with all experiments having a similar duration of 47 hours. (**B**) Top: Rescaled and averaged time series of successive pulse–to– pulse intervals of Class-2 activity (same intervals as in Fig. 2). Bottom: Averaged and rescaled growth rate for the same time intervals, calculated using the same approach as outlined in (A).

**Fig. S8.**
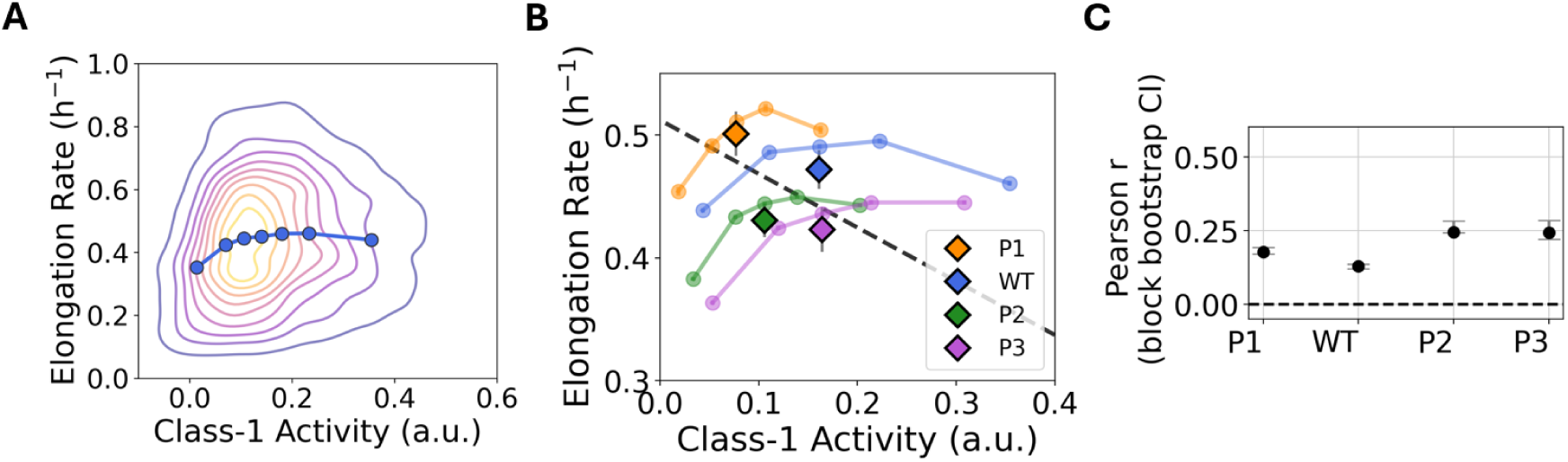
Class-1 activity and elongation rate correlation. (**A**) The color map shows the density of single-cell measurements of elongation rate versus Class-1 activity, for the WT flagellar expression, strain E98KFD. In blue is the mean of the binned data. The dataset includes n=57,592 data points from N=107 mother cell lineages tracked over ∼47 hours. (**B**) Mean binned elongation rate as a function of Class-1 activity, for different strains that have different flagellar expression levels, i.e., strains in which Class-1 activity is controlled by synthetic promoters P1, P2, or P3, and the native promoter, WT. The triangles show the population average for each strain. (**C**) Block-bootstrap estimates of the Pearson correlation between Class-1 activity and elongation rate for each strain, resampling whole lineages to account for within-lineage dependence. Dots show correlations from the original data; vertical lines indicate 95% percentile block-bootstrap confidence intervals.

**Fig. S9.**
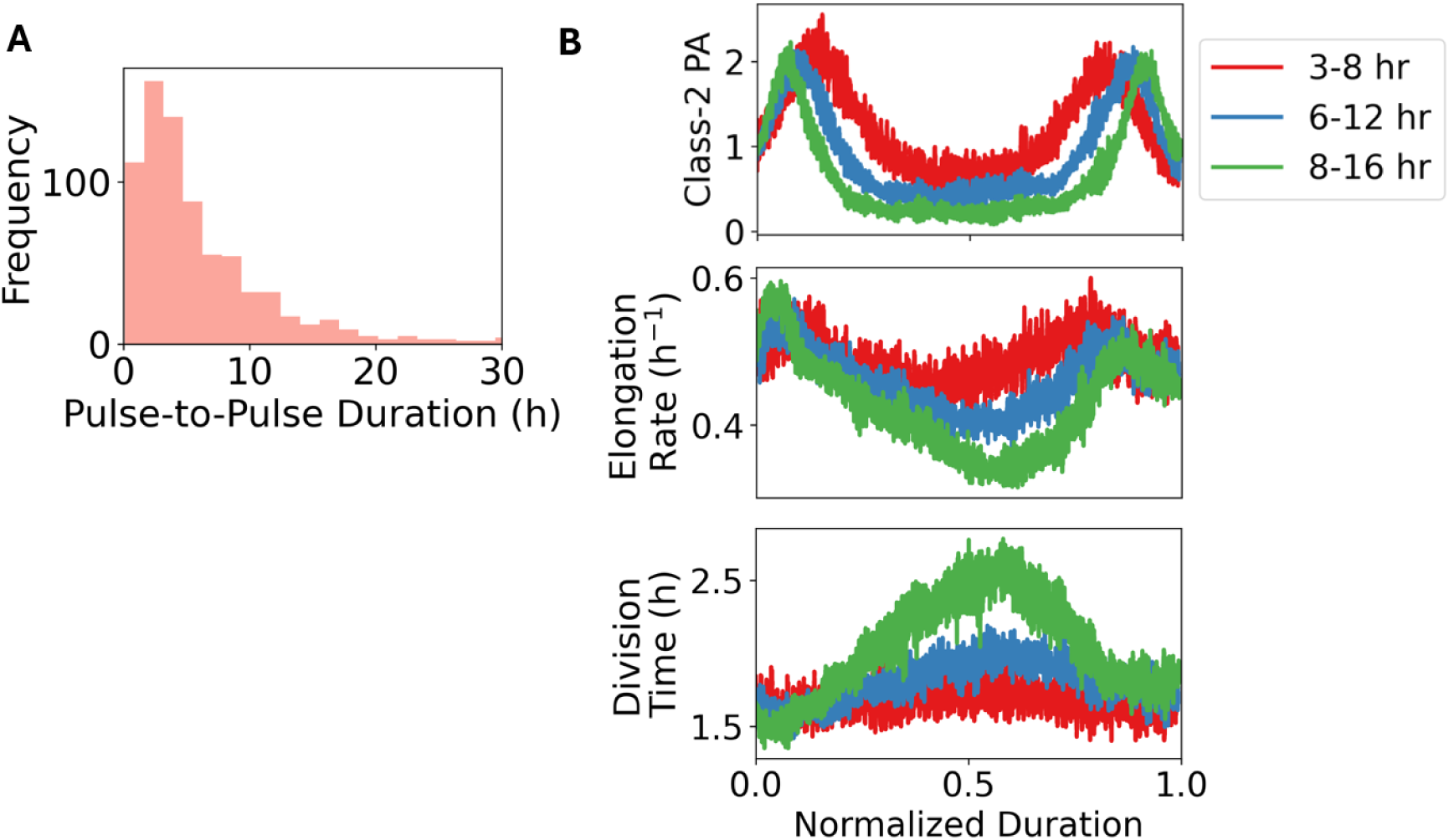
Class-2 pulse-to-pulse and elongation rate analysis. Pulse–to–pulse segments are defined as portions of the Class–2 activity that include two consecutive pulses (*on* – *off* – *on*). (**A**) Histogram of the duration of the segments. (**B**) Average of different variables for pulse-to-pulse intervals of different durations, with red representing 3–8 h, blue representing 6–12 h, and green representing 8–16 h. From top to bottom: Class-2 activity, elongation rate, and division time. The number of intervals (262, 126, 103) corresponds to the time intervals between the two successive pulses ranging from 3-8 h, 6-12 h, and 8-16 h, respectively. Experiment was performed using *E. coli* strain E98KFD in MOPS-Gly medium.

**Fig. S10.**
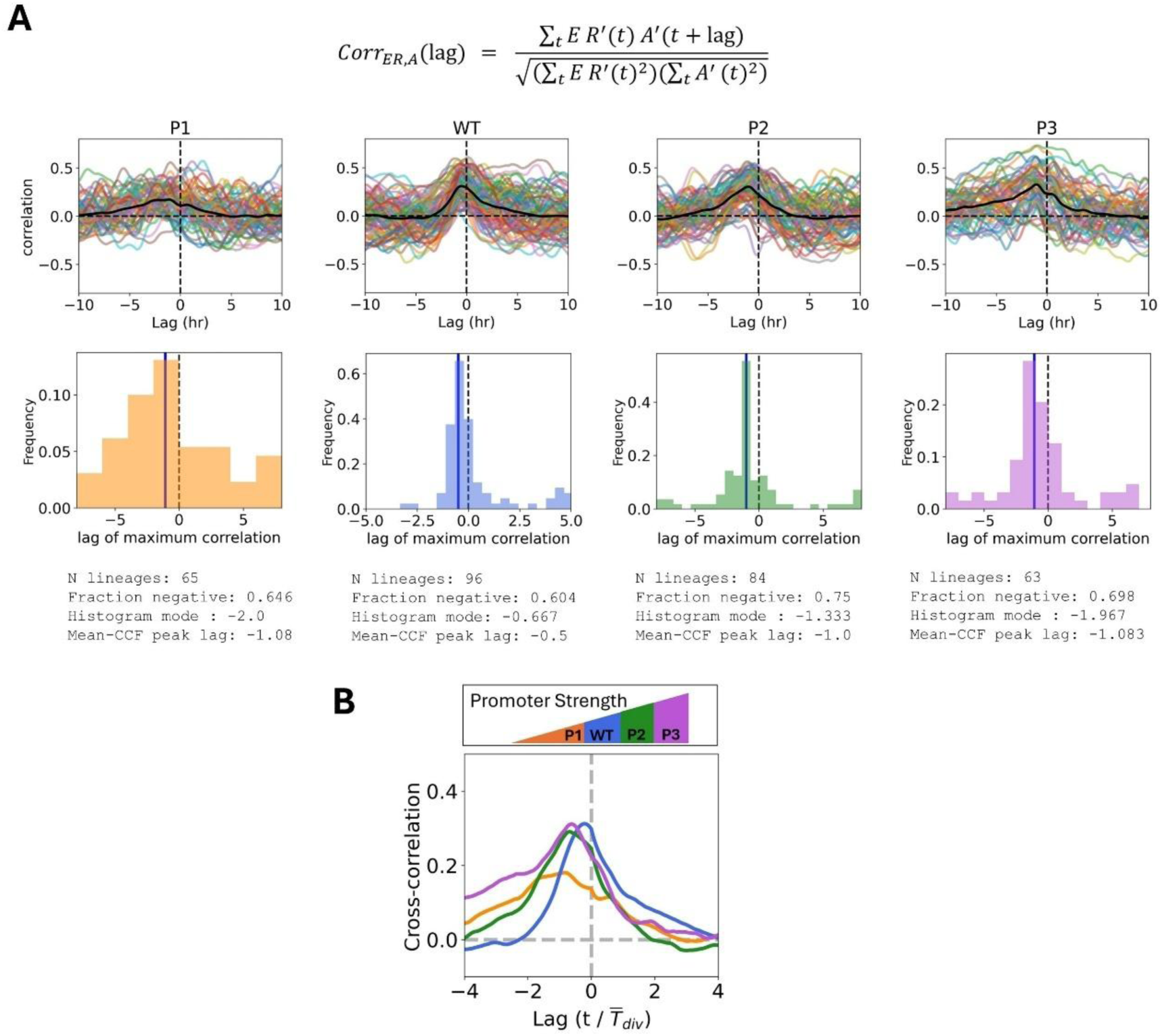
Cross-correlation between elongation rate and Class-2 activity in strains of varying flagellar activity. (**A**) For each flagella-reporter strain, normalized cross-correlation functions between elongation rate and promoter activity were computed separately for every lineage (top, colored lines), using the equation on the top, where *ER*′(*t*) and A’(t) denote the elongation–rate and activity time series, respectively, with their mean subtracted. The cross-correlation quantifies how strongly signals *ER*′(*t*) and A’(t) are related when the activity signal A’(t) is shifted in time (by a lag) relative to the elongation-rate signal *ER*^′^(*t*). These lineage-level cross-correlations were then averaged pointwise across lineages to obtain the mean cross-correlation as a function of lag (top, black line). For each lineage, the temporal delay between elongation rate and promoter activity was defined as the lag within a predefined, biologically relevant window at which the normalized cross-correlation reached its maximum. Positive delays indicate that elongation rate lags behind promoter activity, whereas negative delays indicate that elongation rate precedes promoter activity. The distribution of delays for all lineages of a given strain is shown as a histogram (bottom), where the vertical dotted line indicates zero lag, and the blue line marks the lag at which the mean cross-correlation peaks. (**B**) Mean cross-correlation functions for the different strains, colored as indicated. For each strain, cross-correlations were first computed for individual lineages, and the lag axis was normalized by the average division time (*T_div_*) of that strain before averaging across lineages.

**Fig. S11.**
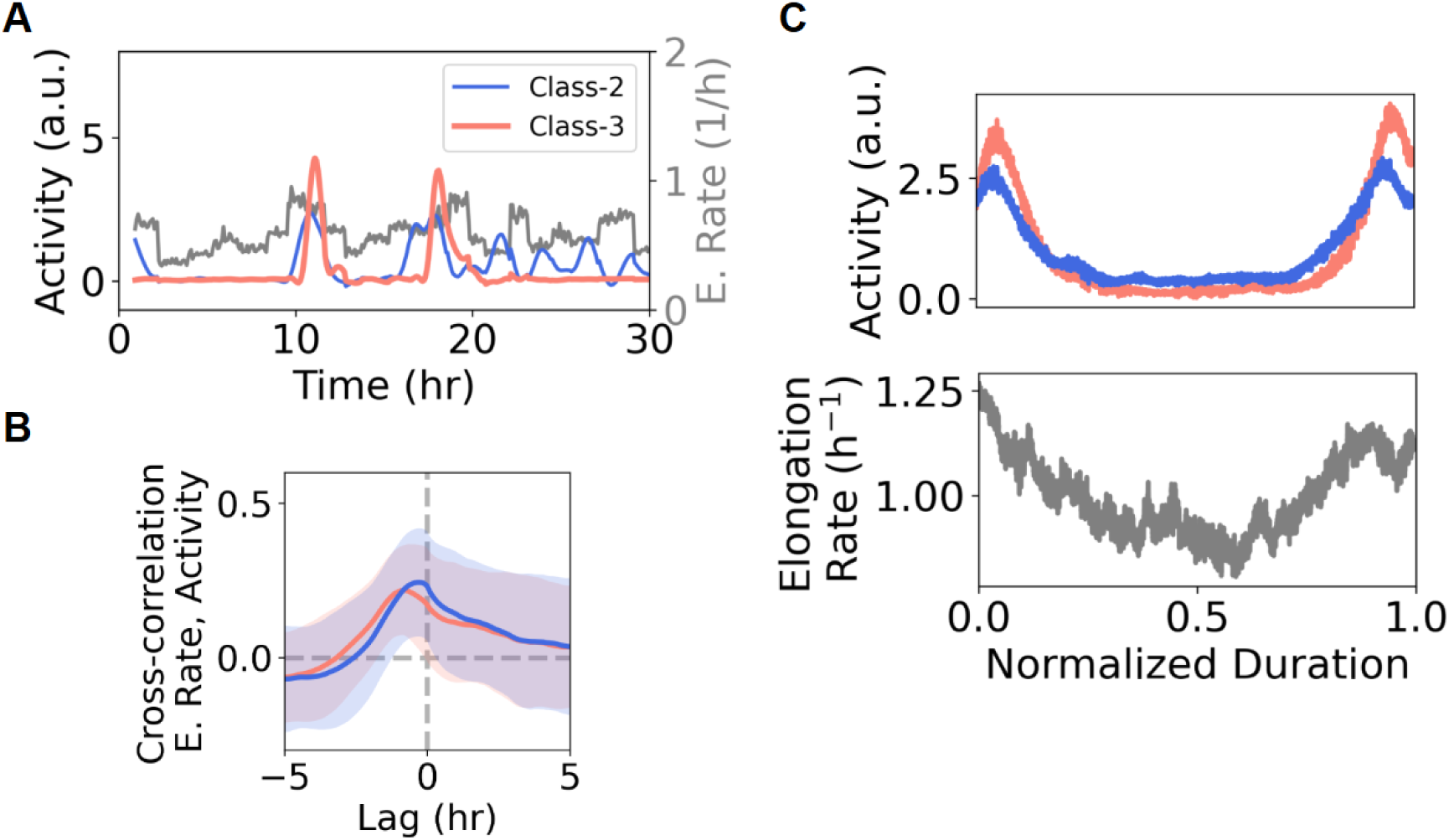
Class-3 activity and elongation rate correlation. (**A**) Time series of flagellar promoter activity for Class-2 (blue; *fliFp*), Class-3 (pink; *fliCp*), and elongation rate (grey) in a representative mother cell across multiple divisions. promoter activity is normalized to its mean over the series. (**B**) Average cross-correlation between Class-2 activity and elongation rate (blue) and between Class-3 activity and elongation rate (pink). Cross-correlation was computed for each lineage (n = 68), and values were averaged across all lineages for each time lag. Shaded areas represent the standard deviation for each lag. The lag corresponding to the maximum (average) cross-correlation is 0.835 h, for Class-3, and 0.245 h for Class-2. (**C**) Following the procedure outlined in Fig. 2, Class-3 activity time series were segmented into intervals spanning two consecutive peaks (on-off-on states). Intervals of similar duration were selected (10-16 h; n = 55 intervals), and their duration was normalized. Activity profiles were then smoothed. For the same intervals defined by Class-3 peaks, we also averaged Class-2 activity and elongation rate (shown below). The bacterial strain used for all these analyses was E98KFC (see Strain List).

**Fig. S12.**
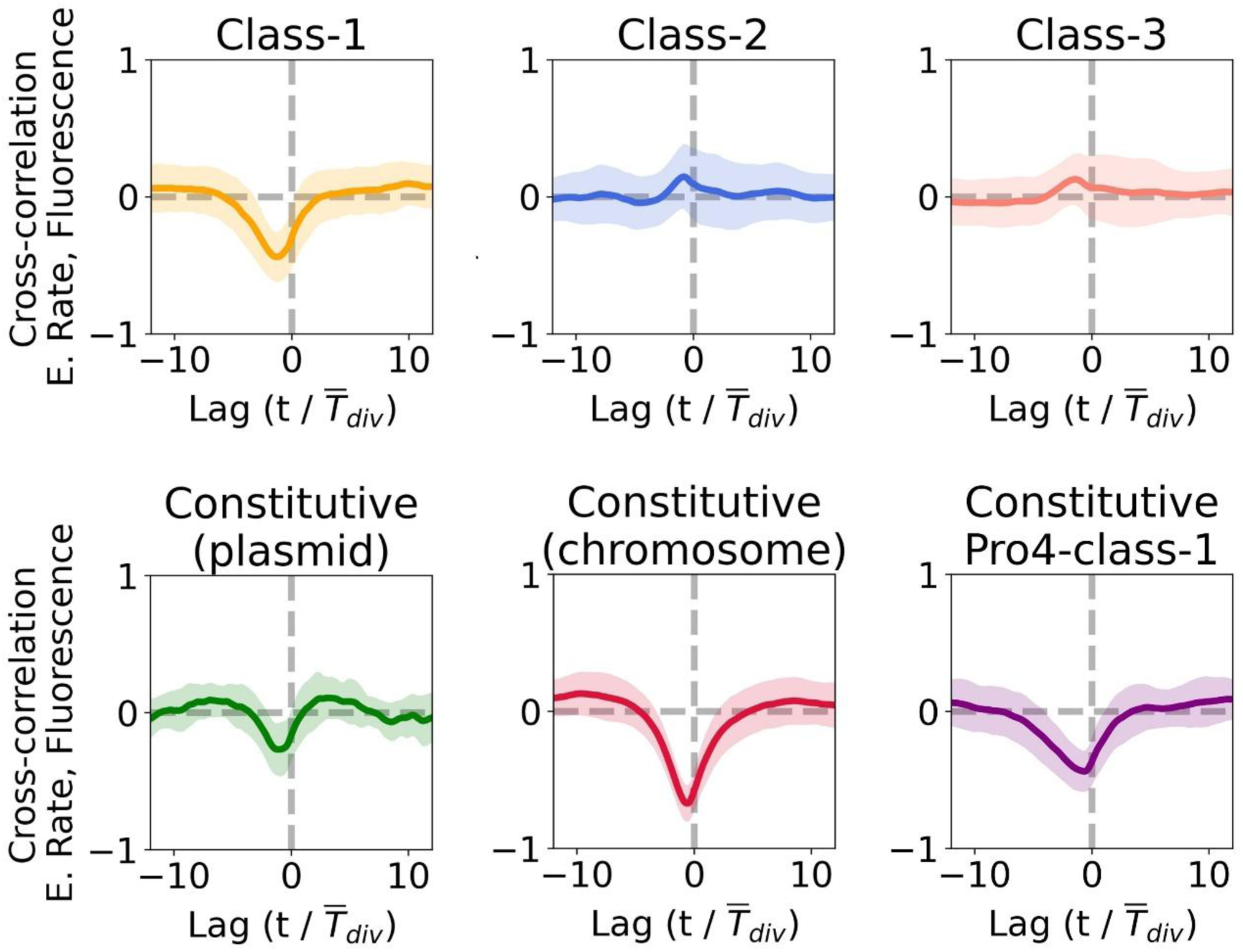
Cross-correlation analysis of elongation rate and fluorescence for various promoters and strains. The first row displays the correlation coefficients over time lag for three promoters, each representing one of the three flagellar classes, controlling the fluorescence: Class-1 (flhDp_mVenusNB; strain E98KFD), Class-2 (fliFp_SCFP3A; strain E98KFD), and Class-3 (*fliCp_mVenusNB*; strain E98KFC). In the second row are the plots corresponding to constitutive expression: Constitutive promoter (Pro4_mVenusNB; strain MGPro4-Venus) encoded on a plasmid, constitutive promoter encoded in the chromosome (pRNAI_mCherry; strain E98KFC), and a constitutive promoter that controls both the Class-1 genes *flhD* and *flhC,* and the gene encoding the fluorescent protein (Pro4-flhDC_mVenusNB; strain E98KFD Pro4_T7). The Fluorescence signal is the average intensity of the cells per area, and the elongation rate is the time derivative of the Size normalized by Size, *ER*(*t*) = (1/*S*(*t*) *dS*(*t*)/*dt*). Both signals were smoothed using a window of 7 frames (35 min). Each plot shows the average of the correlation over many time series, and the shaded region represents the standard deviation about the mean. The fluorescent proteins used were selected for their fast maturation time, as described in [52].

**Fig. S13.**
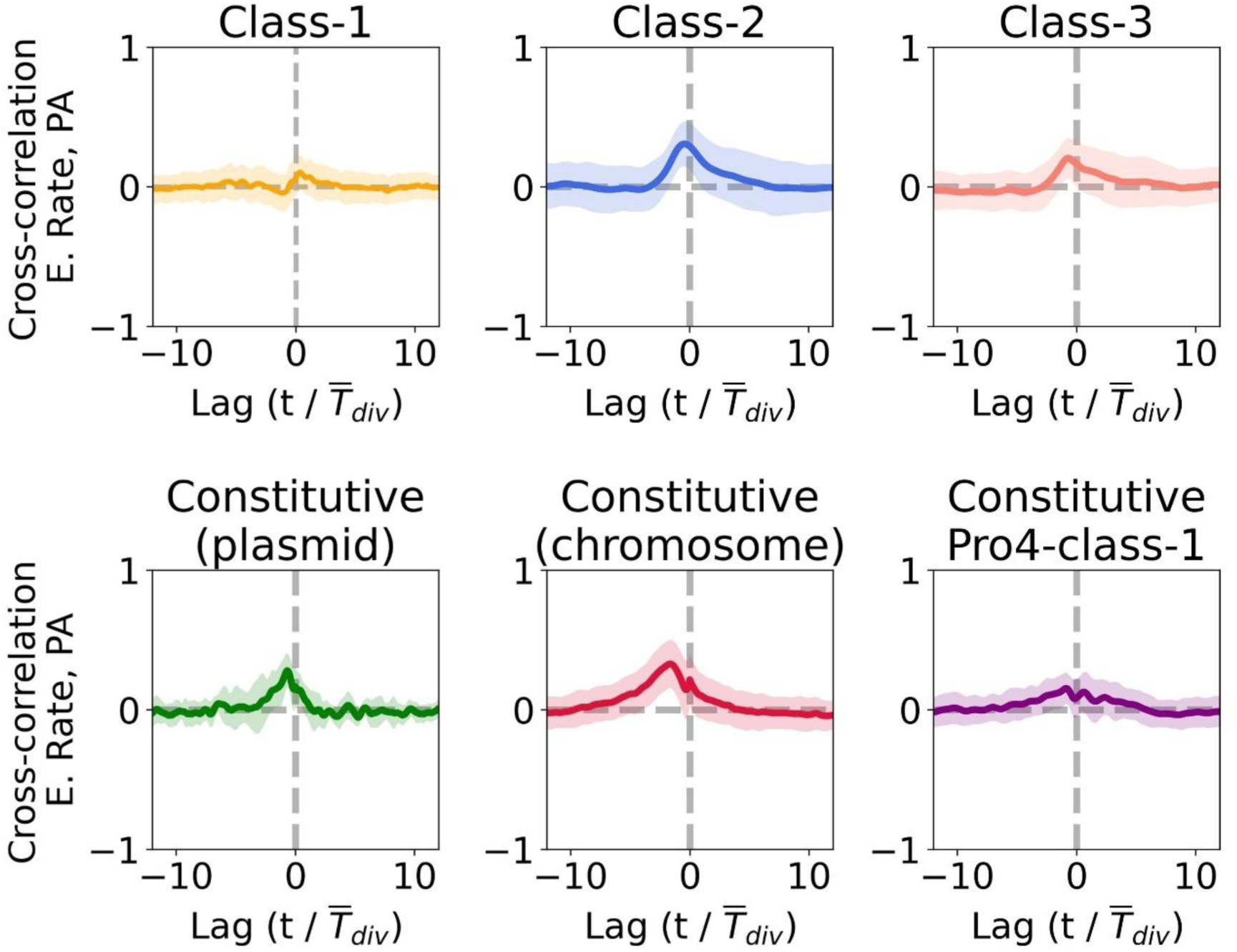
Cross-correlation analysis of elongation rate and promoter activity for various promoters and strains. The promoter activity and elongation rate were computed as described in Materials and Methods. The promoters and strain names appear in the same order as in Fig. S12. Sampaio et al. [22] monitored the correlation between growth rates and reporter expression levels from various *E. coli* promoters at the single-cell level, and found anticorrelation. When we calculated the cross-correlation between the elongation rate and reporter levels (mean fluorescence), instead of promoter activity, we similarly observed a negative correlation for reporters driven by constitutive promoters (Class-1 and synthetic types; Fig. S12). However, for reporters driven by Class-2 and Class-3 promoters, we observed a positive correlation between the growth rate and both reporter levels and promoter activity.

**Fig. S14.**
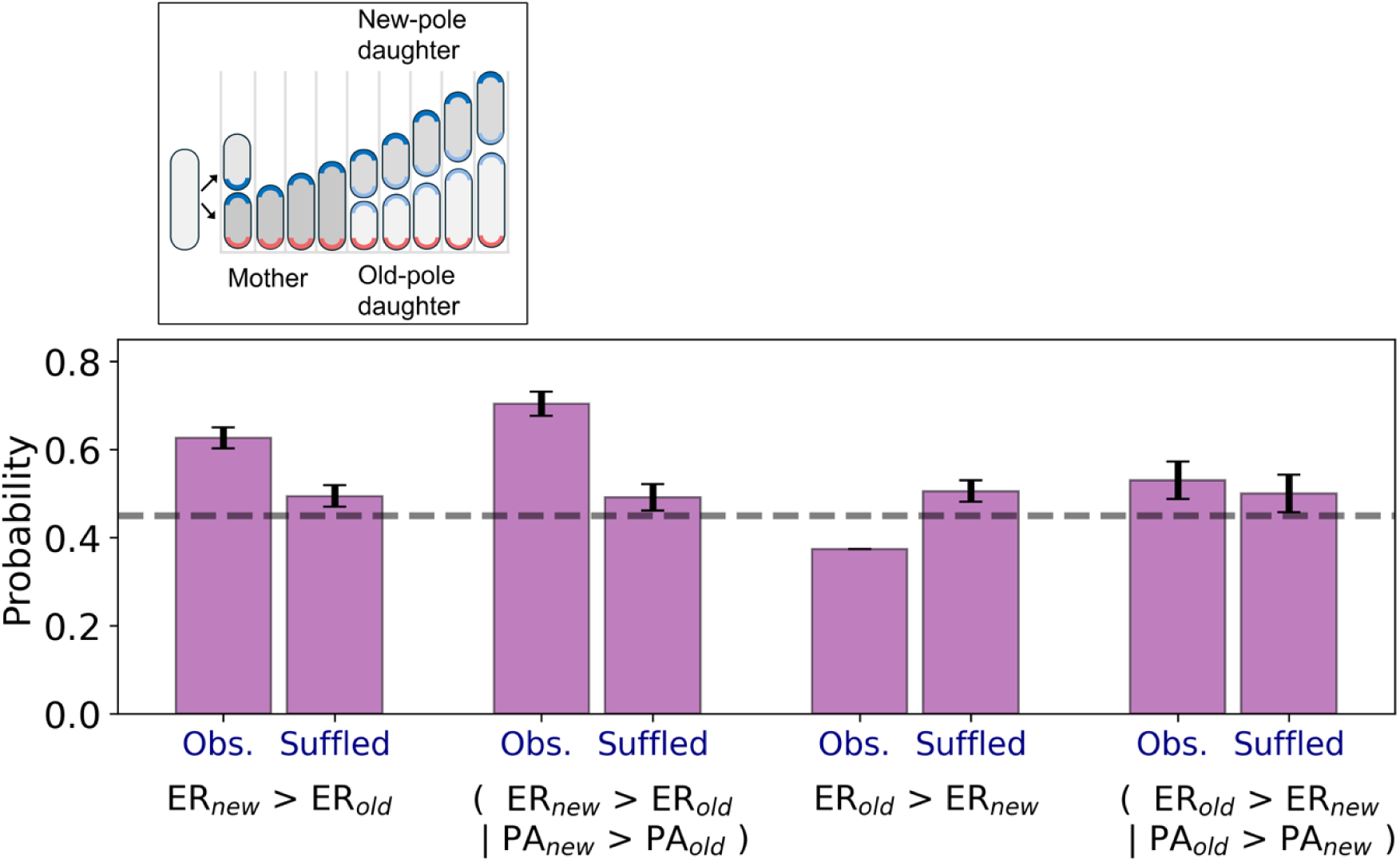
Probability of bias advantage between the elongation rate of old and new pole daughters. Probability that old pole daughters (newly born cells at the close end of the channel which inherits the old pole (red pole in the diagram)) or new pole daughters (daughter cells born closer to the open side of the channels that inherits the mother’s new pole (dark blue)) have higher elongation rates, with and without conditioning on promoter activity. Bars show the probability that the daughter with the elongation rate of the new pole (*ER_new_*) has a higher elongation rate than the old daughter (*ER_new_*> *ER_old_*), or vice versa (*ER_old_*> *ER_new_*), either unconditionally or conditional on that daughter having higher promoter activity than its sister (e.g. *ER_new_* > *ER_old_* given *PA_new_* > *PA_old_*). For each case is shown the probabilities of the observed data, ‘Obs.’, and the ‘Shuffled’ pairs’ bars show the corresponding probabilities under a within– pair permutation null in which elongation rates are randomly swapped between sisters with probability 0.5. Error bars indicate 95% Clopper–Pearson exact binomial confidence intervals. The horizontal dotted line indicates a probability of 0.5, corresponding to no bias between sisters.

**Fig. S15.**
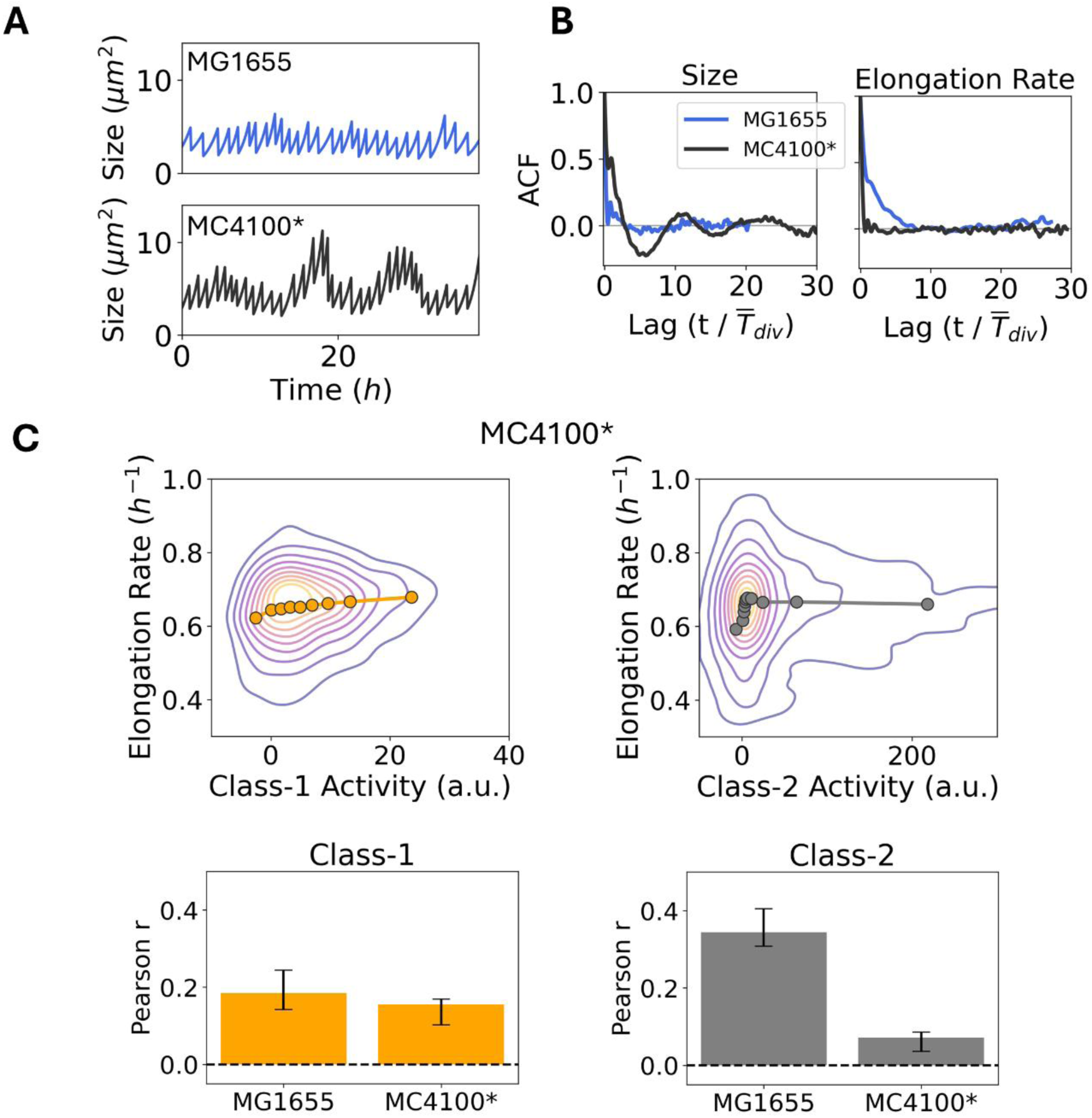
MG1655 and MC4100 differ in size-oscillation dynamics but not in short-timescale growth– expression coupling, which is attenuated only at flagellar Class-2 activity. (**A**) Typical time series showing the Size of MG1655 background strain (EMGFD) and MC4100* (MC4100 variant with flagellar expression but non-motile). (**B**) Autocorrelation function (ACF) of the Size time and Elongation Rate series in MG1655 (blue) and long-term oscillating MC4100* strain (black). Averages of individual ACFs were calculated from ∼60 lineages and normalized by the mean division time. (**C**) Top: Density of single-cell measurements of elongation rate versus promoter activity for Class-1 and Class-2 promoters in MC4100*. Circles show the mean of the binned data. Bottom: Pearson correlation coefficients between elongation rate and promoter activity for Class-1 and Class-2 in MG1655 and MC4100, estimated by block bootstrap to account for temporal autocorrelation within time series; error bars denote 95% confidence intervals.

**Fig. S16.**
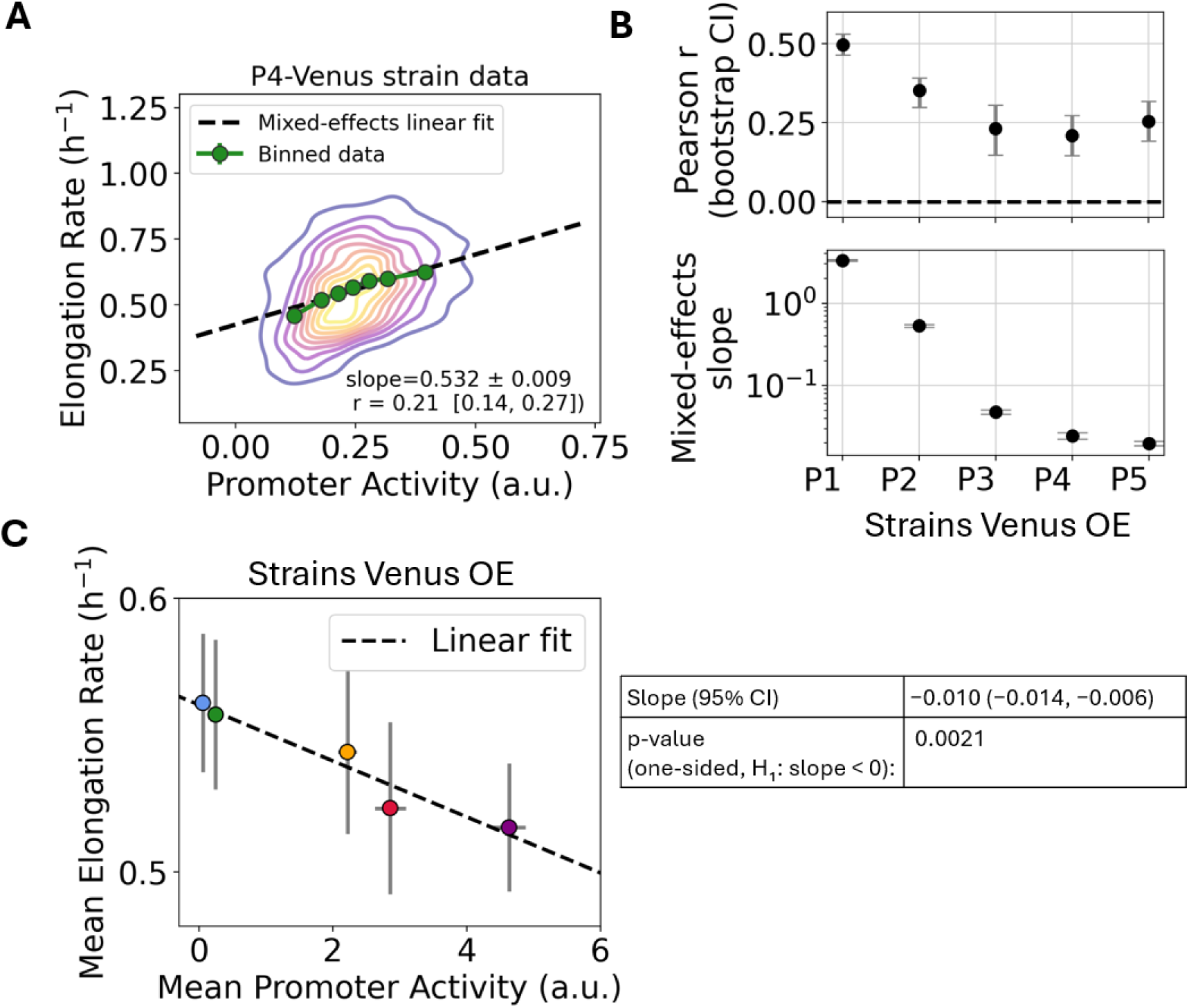
Statistical analysis of activity–elongation rate relationships in constitutive fluorescent reporter strains. **(A)** The color map shows the density of single-cell measurements of elongation rate versus promoter activity, using the P4-Venus strain. In green is the mean of the binned data. The black dashed line shows the fixed effect prediction from the mixed-effects linear model (ER ∼ PA with random intercepts per lineage), representing the average single-cell trend after accounting for lineage-to-lineage variability. **(B) Top:** Block bootstrap estimates of the Pearson correlation between activity and elongation rate for the strains that overexpress Venus, resampling whole lineages to account for within lineage dependence. Dots show correlations from the original data; vertical lines indicate 95% percentile block bootstrap confidence intervals. **Bottom**: Mixed-effects linear regression of elongation rate on promoter activity for each strain. Each point shows the fixed-effect slope (change in ER per unit PA); horizontal lines indicate 95% confidence intervals from a mixed model with random intercepts per lineage. (**C**) Mean Elongation rate as a function of the mean promoter activity, the error bars show the standard error of the mean and the dotted line shows the weighted linear regression fit (weights proportional to the number of lineages per strain), illustrating the population-level trend across strains. The table summarizes the weighted linear regression of strain averaged ER on strain-averaged PA, showing the negative slope, its 95% confidence interval, and the one-sided p-value.

**Fig. S17.**
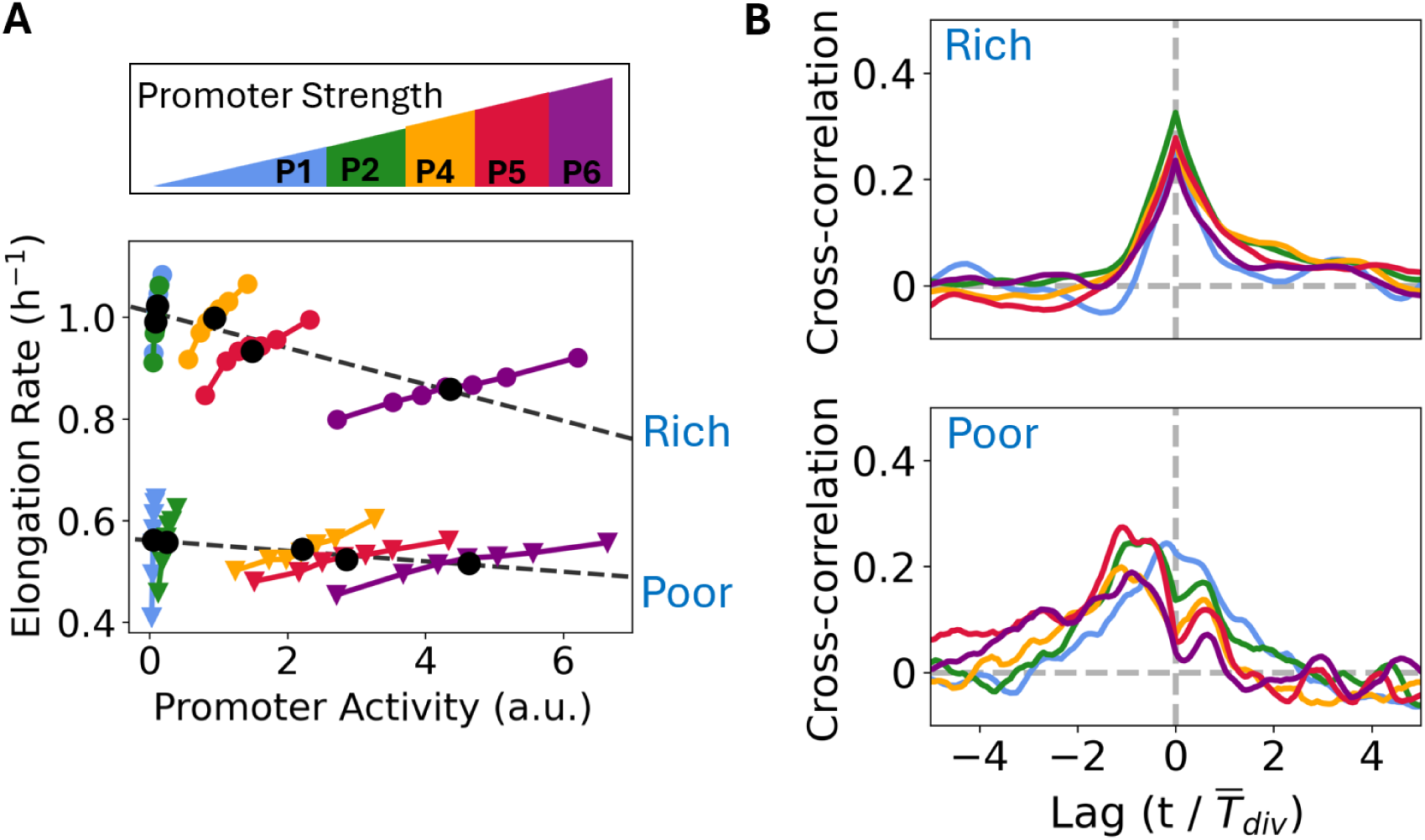
Elongation rate and promoter activity across constitutive expression levels and media. (**A**) For a set of strains expressing the fluorescent protein Venus, driven by promoters of varying strengths (P1 to P6, as indicated at the top), and grown in two different media (labeled Rich and Poor), we calculated the mean elongation rate as a function of binned promoter activity. The black dots represent the population average for each strain, while the lines indicate the linear fit of these points for both conditions (cells grown in either rich or poor medium). (**B**) The average cross-correlation (see Materials and Methods for the definition) between elongation rate and promoter activity was calculated for the lineages of strains producing unnecessary proteins (same data and color-coding as in panel A), for both poor and rich media. Binning and cross-correlation analyses were performed on time-series of approximately 37 hours in duration, obtained from n = (30, 21, 19, 14, 24) lineages under poor medium and n = (12, 33, 39, 22, 40) lineages under rich medium, corresponding to strains carrying promoters P1 through P6, respectively. Poor medium consisted of Teknova MOPS EZ defined base (no ACGU) supplemented with 0.4% glycerol, 1.32 mM phosphate, and 0.85 g/L Pluronic F-108. Rich medium comprised the same MOPS base with Supplement EZ, ACGU solution, 0.5% glucose, 1.32 mM phosphate, and 0.85 g/L Pluronic F-108.

**Fig. S18.**
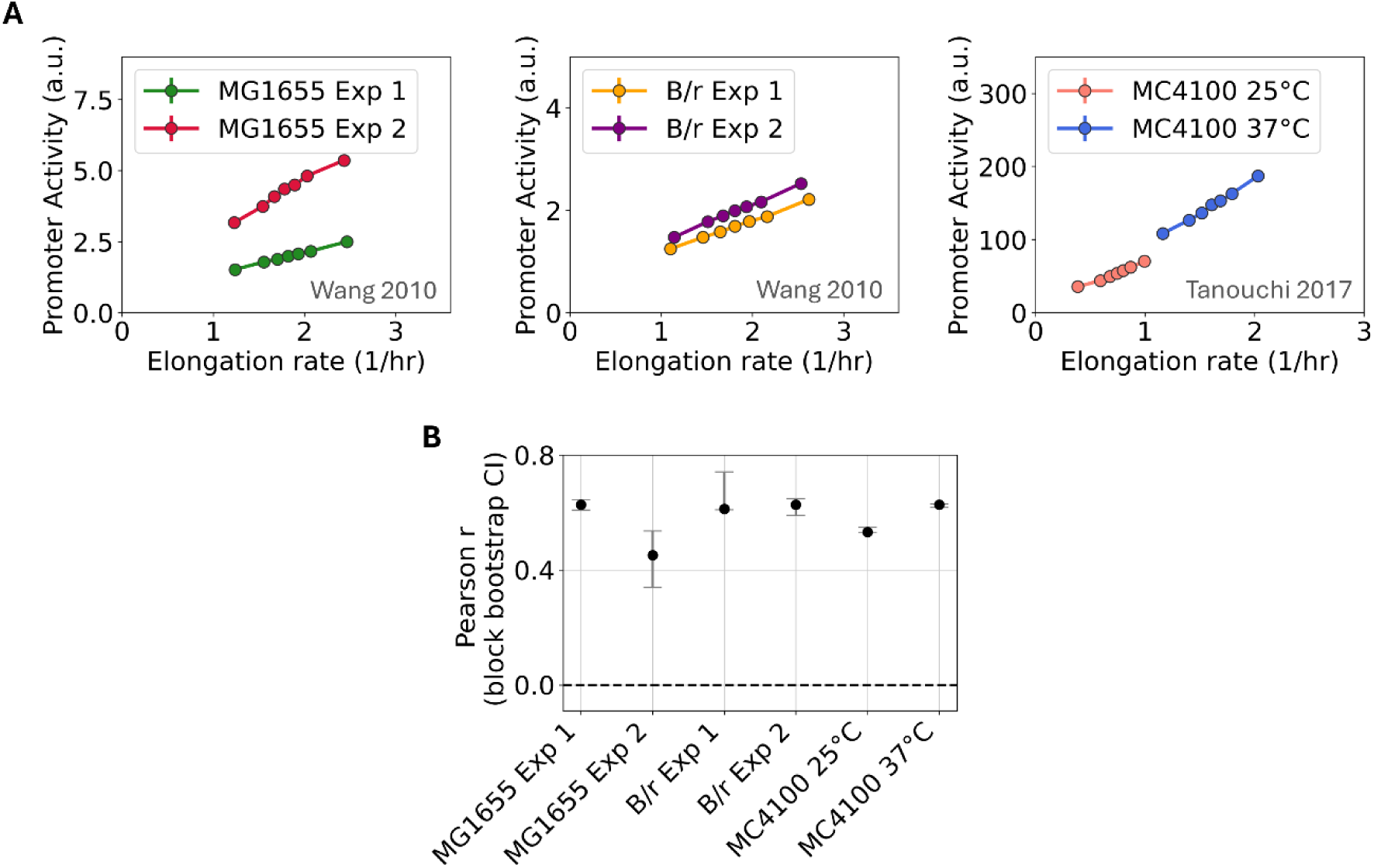
Growth-constitutive expression coupling is conserved across *E. coli* strains and conditions. (**A**) Density of single-cell measurements of elongation rate versus promoter activity for constitutively expressed fluorescent protein in MG1655 (2 experiments, N = 508, 201 cells), B/r (2 experiments, N = 231, 932 cells), and MC4100 at 25°C and 37°C (N = 669, 259 cells). Data for MG1655 and B/r are from Wang et al. [33]; data for MC4100 are from Tanouchi et al. [44, 48]. We selected time traces for reanalysis after excluding those affected by segmentation errors. Circles show the mean of binned data. (**B**) Pearson correlation coefficients between elongation rate and promoter activity for each strain/condition shown in (A), estimated by block bootstrap to account for temporal autocorrelation within time series. Error bars denote 95% confidence intervals.

**Table S1.**
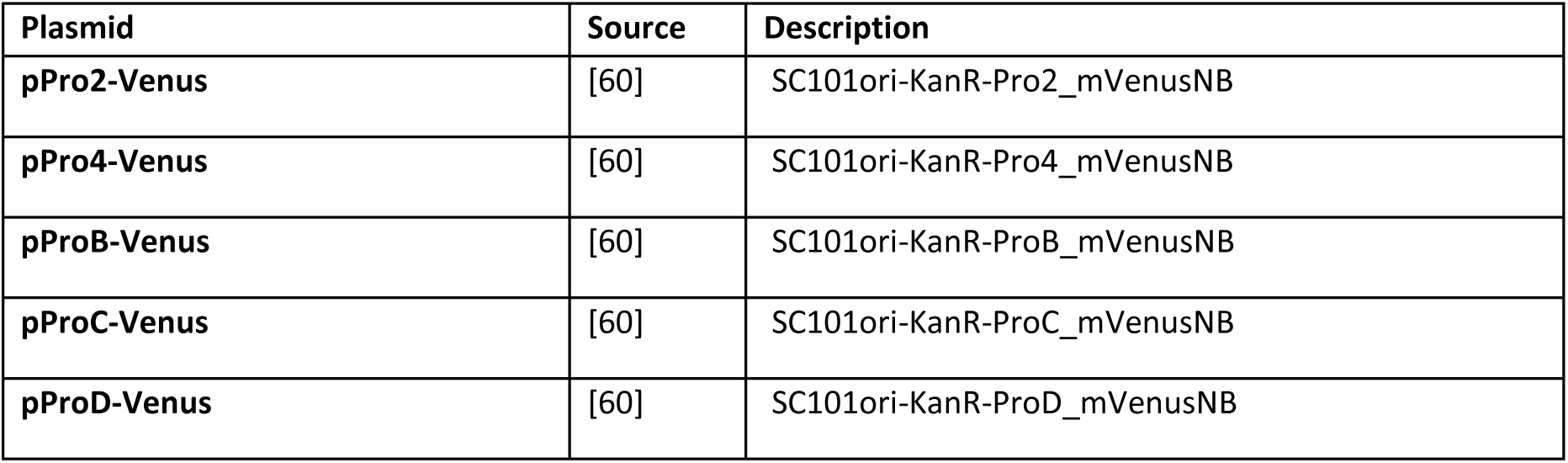
List of plasmids.

| Plasmid | Source | Description |
| --- | --- | --- |
| pPro2-Venus | [60] | SC101ori-KanR-Pro2_mVenusNB |
| pPro4-Venus | [60] | SC101ori-KanR-Pro4_mVenusNB |
| pProB-Venus | [60] | SC101ori-KanR-ProB_mVenusNB |
| pProC-Venus | [60] | SC101ori-KanR-ProC_mVenusNB |
| pProD-Venus | [60] | SC101ori-KanR-ProD_mVenusNB |

**Table S2.**
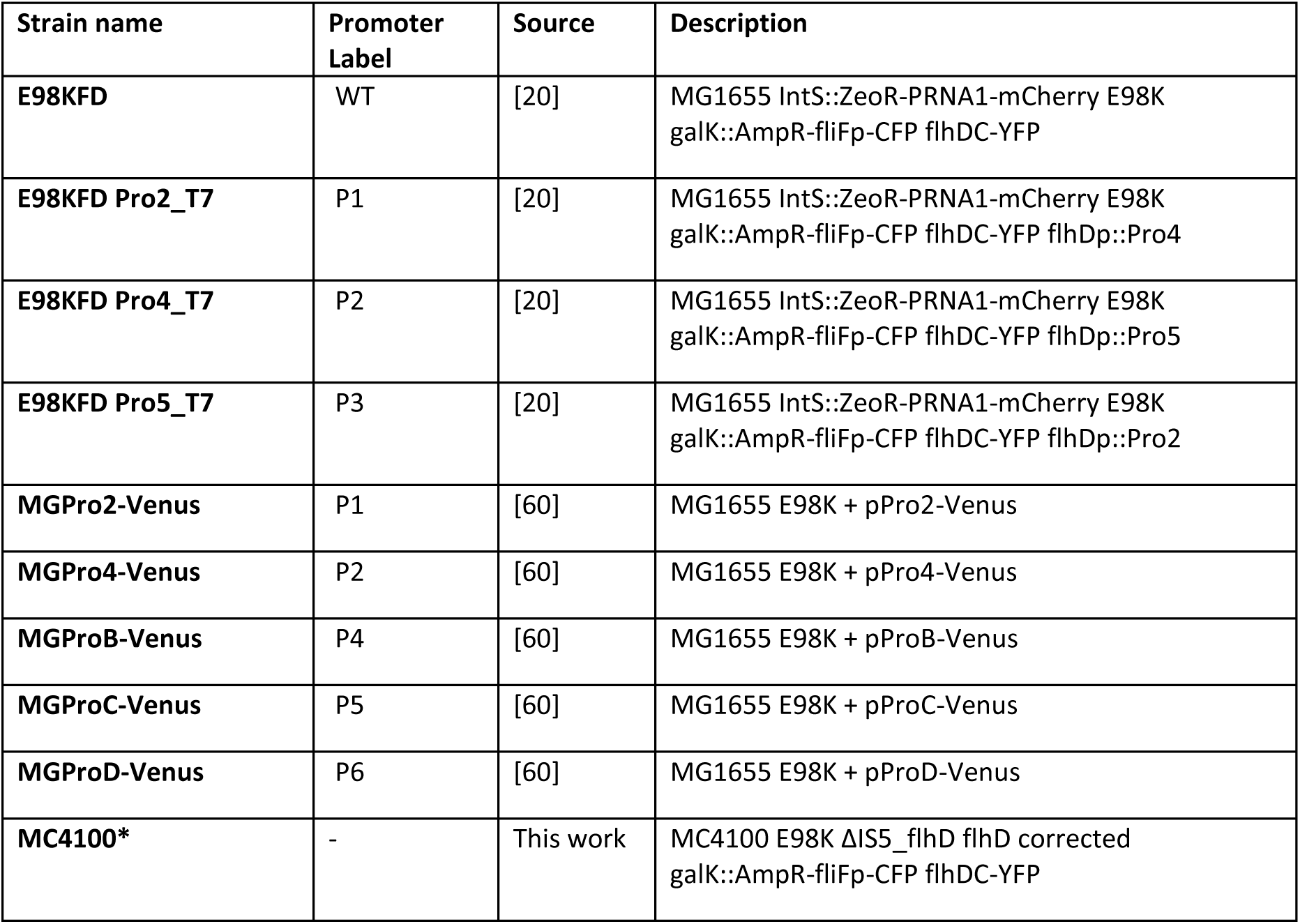
List of strains.

**Table S3.** Estimated mVenus proteome fraction across promoter strains.

| Strain | Promoter name | Average Elongation Rate ($h^{-1}$) | % reduction vs. WT | Estimation $\Phi_U$ (% dry cell weight) |
| --- | --- | --- | --- | --- |
| MGPro2-Venus | P1 | 0.5618 | 1.29% | 0.55% |
| MGPro4-Venus | P2 | 0.5575 | 2.04% | 0.87% |
| MGProB-Venus | P4 | 0.5438 | 4.44% | 1.91% |
| MGProC-Venus | P5 | 0.5232 | 8.05% | 3.46% |
| MGProD-Venus | P6 | 0.51621 | 9.30% | 3.99% |

'% reduction' is calculated relative to the WT (non-expressing) growth rate $\mu_{max} = 0.5691 h^{-1}$ . $\Phi_U$ is estimated using the ribosome-allocation relationship $\mu/\mu_{max} = 1 - b \cdot \Phi_U$ (Scott et al., [6]), with $b = 2.33 \mu g$ DCW per $\mu g$ protein, the empirically determined value for GFP-family proteins (Bienick et al., 2014 [47]).

